# Identification of target genes and regulatory networks for bone mineral density GWAS loci through systematic targeting and inhibition

**DOI:** 10.64898/2026.02.11.705362

**Authors:** Mohammadjavad Kahlani, Larry Mesner, Vedavathi Madhu, Lucia Wagner, Kristyna Kupkova, Hannah J. Pitas, Marinella Gebbia, Mehrdad Kazemi, Chiraz Belarbi, Ana C. Mello, Samuel Ghatan, Tuuli Lappalainen, Neville E. Sanjana, John A. Morris, Charles R. Farber

## Abstract

Osteoporosis is characterized by low bone mineral density (BMD) and elevated fracture risk. Most BMD-associated GWAS variants lie in noncoding regions, complicating efforts to identify causal genes and mechanisms. To overcome this variant-to-function challenge, we previously developed STING-seq, a framework integrating biobank-scale GWAS with single-cell CRISPR inhibition (CRISPRi) screening to directly connect noncoding *cis*-regulatory elements (CREs) to their target genes. Applied to human fetal osteoblast (hFOB) cells across osteogenic differentiation, STING-seq linked 76 CREs to 75 target genes at BMD loci. Arrayed CRISPRi and activation validated key CRE–gene relationships, including long-range enhancers regulating *CXCL12* (over 500 kb apart). CRISPRi effects were broadly reproduced in primary human mesenchymal stem cells, with 82% of regulatory effects concordant in direction with hFOB cells and strongly correlated in magnitude (Pearson r = 0.83). We further uncovered *trans*-regulatory networks and characterized the *DAP3*–*YY1AP1* bidirectional promoter, demonstrating a role for *DAP3* in mineralization via mitochondrial pathways. Together, these findings provide mechanistic insight into how noncoding GWAS variants shape osteoblast activity and highlight the genes and pathways that mediate genetic effects on BMD.

## Introduction

Osteoporosis is a highly prevalent bone disorder characterized by low bone mineral density (BMD) and increased fracture risk ^1^. Genome-wide association studies (GWAS) have identified over 500 loci associated with BMD, offering valuable insights into the genetic architecture underlying osteoporosis ^2–5^. However, translating GWAS findings to actionable genetic targets is challenging due to the majority of GWAS-identified genetic variants mapping to noncoding regions. This variant-to-function problem makes it difficult to pinpoint causal variants and identify the specific genes these variants influence.

The variant-to-function problem concerns the challenge of determining how genetic variants, particularly those in noncoding regions, influence genes, cellular processes, and human phenotypes ^6–15^. More than 90% of GWAS-associated variants are noncoding ^16^, and resolving their functions requires approaches that connect statistical associations to regulatory elements, target genes, and ultimately cellular phenotypes. Sequence-based models such as Enformer ^17^, Borzoi ^18^, and AlphaGenome ^11^ can predict diverse regulatory features from genomic sequence, but reliably identifying the target genes and cellular consequences of noncoding variants remains challenging. Complementary approaches therefore integrate GWAS and cell-type-relevant epigenomic data to prioritize regulatory elements and predict enhancer–gene relationships. For example, the Activity-by-Contact model uses enhancer activity and chromatin contacts to predict regulatory connections ^19,20^, while related models such as ENCODE-rE2G integrate perturbation data to improve enhancer–gene prediction ^21^.

CRISPR-based functional genomics provides an experimental framework for testing these predictions. Perturb-seq has enabled systematic validation of enhancer–gene connections, coupling genetic perturbations with single-cell transcriptomic readouts to map gene function and downstream cellular programs ^6,22–24^. We previously developed Systematic Targeting and Inhibition of Noncoding GWAS loci with single-cell sequencing (STING-seq), which combines GWAS-guided *cis*-regulatory element (CRE) selection, pooled CRISPRi, and single-cell RNA sequencing to experimentally connect variant-associated CREs to their target genes and downstream transcriptional programs ^6^. Recently, GWAS-informed noncoding CRISPRi screens have begun to apply this framework to BMD-associated loci, illustrating its potential to identify effector genes in osteoblasts ^25^.

Noncoding GWAS variants can influence cellular phenotypes at multiple regulatory levels. Locally, a variant may alter the activity of a CRE and thereby affect one or more target genes in *cis*. These effects may then propagate in *trans* through broader regulatory networks, particularly when the target gene encodes a transcriptional or cellular regulator. Resolving GWAS loci therefore requires both identification of the immediate target genes of variant-associated CREs and characterization of their downstream effects on cellular programs and phenotypes. Here, we applied STING-seq in human fetal osteoblast (hFOB) cells across three stages of osteogenic differentiation, with targeted replication in primary human mesenchymal stem cells (hMSCs), to identify target genes of noncoding BMD GWAS loci. We linked 76 CREs to 75 target genes and validated select regulatory relationships using CRISPRi and CRISPR activation (CRISPRa). We further examined their effects on mineralization and identified long-range enhancers regulating established BMD genes, including *CXCL12*, as well as previously unrecognized roles for *SMARCD3* in osteoblast differentiation and *DAP3* in mitochondrial function and mineralization. Beyond these *cis*-regulatory relationships, we identified downstream regulatory networks, including a *TRIP11*-associated network enriched for genes underlying monogenic skeletal disorders, suggesting that regulatory perturbations at BMD loci can converge in *trans* on established determinants of skeletal biology. Together, these findings demonstrate how STING-seq can connect noncoding GWAS loci to effector genes, regulatory networks, and disease-relevant cellular phenotypes.

## Results

### A validated human bone-forming cell model

Although many cell types both intrinsic and extrinsic to bone influence BMD ^26^, several recent studies have shown that *cis*-regulatory regions and genomic intervals flanking highly expressed genes in osteoblast lineage cells are strongly enriched for BMD heritability ^2,25,26^, underscoring their key role in mediating genetic influences on bone. To perform STING-seq in osteoblasts, we selected the hFOB cell line, a well-established hMSC line capable of osteogenic differentiation. The hFOB cell line was derived from female fetal bone tissue and in addition to osteogenic differentiation, it is also capable of chondrogenic and adipogenic differentiation, akin to primary hMSCs ^27,28^. hFOB cells express surface markers of hMSCs including CD90, CD44, CD73, and CD105, and are negative for the hematopoietic markers CD34 and CD45 ^28^. Moreover, hFOB cells are karyotypically stable and do not exhibit tumorigenic properties, distinguishing them from commonly used osteoblast-like cell lines derived from osteosarcomas ^28^. Lastly, hFOB cells are capable of generating bone when transplanted into mice ^29^.

We previously performed short-read ^30^ and long-read ^31^ RNA sequencing of hFOB cells at days 0, 2, 4, and 10 of osteogenic differentiation (**Methods**). To evaluate the extent to which hFOB cells recapitulate osteoblast differentiation *in vivo*, we compared the expression of established skeletal lineage markers in a single-cell atlas of the human bone marrow niche (**Extended Data Fig. 1a**) ^26^ with expression across hFOB differentiation time points (**Extended Data Fig. 1b–c, Supplementary Table 1a**). hFOB cells expressed genes associated with mesenchymal stromal and the osteoblast lineage, including *RUNX2*, *DLX5*, *SATB2*, *COL1A1*, *COL1A2*, and *SPARC*, with expression patterns consistent with progression from an osteoblast progenitor-like state from days 0, 2, and 4 to a more mature osteoblast phenotype by day 10. We did not observe markers of osteocytes (*DMP1*, *MEPE*, *SOST*, and *FGF23*). These results suggest that hFOB osteogenic differentiation recapitulates early-to-mid osteoblast maturation.

To identify regions of candidate CREs (cCREs) harboring BMD GWAS variants, we performed ATAC-seq ^32^ at the same time points (four replicates per time point) (**Extended Data Fig. 2a**, **Methods**). We detected an increasing number of ATAC-seq peaks per time point, observing a gain of approximately 40,000 peaks at each of the two-day intervals after osteogenic induction (days 2 and 4), before a reduction of approximately 90,000 peaks by day 10 (**Extended Data Fig. 2b**, **Supplementary Table 1b)**. Approximately 100,000 peaks are shared across all four time points, with linkage disequilibrium score regression (LDSC) analyses revealing that hFOB ATAC-seq peaks captured between 24.6-28.6% of BMD trait heritability (*p*_LDSC_ < 1.2 × 10^-11^) across the four time points (**Supplementary Table 1b**).

Next, we engineered hFOB cell lines for CRISPRi, for discovery screening, and CRISPRa, for validation experiments. For CRISPRi, we previously used a dual-effector vector, KRAB-dCas9-MeCP2, as we and others found it outperformed the single-effector (KRAB-dCas9) system, when targeting CREs, with an effect radius of less than 1 kilobase (kb) ^6,33^. We therefore engineered hFOB cells to constitutively express this dual-effector CRISPRi vector ^6,33^ (**Methods**). For CRISPRa, we used the dCas9-VPR system ^34^, as we previously found it produced robust transcriptional activation at sites of accessible chromatin ^35^. Lastly, we validated that hFOB CRISPRi and CRISPRa cells maintained the potential to mineralize in vitro (**Methods**), enabling downstream analyses to determine CRE or gene effects on osteoblast activity (**Extended Data Fig. 3)**.

### Intersecting BMD GWAS variants with hFOB open chromatin to identify cCREs

We previously performed a GWAS of BMD estimated from quantitative heel ultrasounds and identified 518 loci that explained up to 20.3% of its variance ^2^. We performed statistical fine-mapping to identify likely causal variants ^36^ and constructed 95% credible sets for 499 loci. We required variants to have a posterior inclusion probability (PIP) of at least 1%, nominating 17,003 potentially causal variants (**Supplementary Table 2a**). Of these, 716 were protein coding and 155 were predicted to have deleterious effects on protein function. Of these loci, 109 had a credible set where a protein coding variant had a predicted impact, directly identifying the likely causal gene (**Supplementary Table 2a**).

We previously observed that CREs for transcription factors (TFs) can impact large gene regulatory networks ^6^. As downstream *trans*-regulatory effects are highly informative for understanding the disease mechanisms of *cis*-regulatory targets, we designed guide RNAs (gRNAs) to target cCREs predicted to regulate TFs and thereby uncover novel gene regulatory networks downstream of BMD GWAS loci in osteoblasts. We applied our variant prioritization workflow to (i) hFOB cCREs harboring noncoding fine-mapped variants for which one of the three nearest genes was a TF and (ii) TFs directly nominated by protein coding fine-mapped variants (**Fig. 1a**). Therefore, we intersected fine-mapped noncoding BMD GWAS variants with hFOB ATAC-seq peaks and prioritized variants if one of the three closest expressed gene transcription start sites (TSS) corresponded to a gene annotated as a potential or known TF ^37^, given that most often the closest gene is the target gene ^6,38^. This resulted in a list of 570 fine-mapped noncoding variants that when collapsed into discrete 1 kb windows for CRISPRi, due to its 1 kb effect radius ^6^, produced a list of 393 variants. We then designed three gRNAs per variant and generated off-target Hsu-Scott scores ^39^, removing gRNAs with predicted off-targets. This resulted in a list of 361 variants, corresponding to 138 loci, targeted by three gRNAs each (**Fig. 1b, Supplementary Table 2b**). We also included TSS-targeting gRNAs for 18 TFs, or predicted TFs, identified from protein coding fine-mapped BMD variants. We previously found that fine-mapped protein coding variants can directly nominate novel genes for bone formation with causal effects on mineralization ^2,3^, therefore these 18 genes represent the most likely target genes at their respective loci (**Fig. 1b**, **Supplementary Table 2c**). Some genes had multiple TSSs identified from hFOB long-read RNA-seq, therefore we targeted each TSS with three gRNAs each if they were further than 1 kb apart. Recent studies have also highlighted that CRISPRi-specific off-target effects can occur when a PAM (NGG for SpCas9) occurs in the 12-nucleotide seed region of the 20-nucleotide spacer sequence ^40–42^. We computed seed scores for each guide in our library (**Supplementary Table 2d**). We also included 30 non-targeting gRNAs (no match in the human genome) ^43^ to evaluate differential expression calibration, and 22 *CD55* targeting gRNAs to perform a high multiplicity-of-infection (MOI) titration (**Supplementary Table 2d**). A high MOI experiment allowed us to increase statistical power to detect significant changes in gene expression with fewer cells, as we and others have demonstrated ^6,23,44,45^.

**Figure 1.**
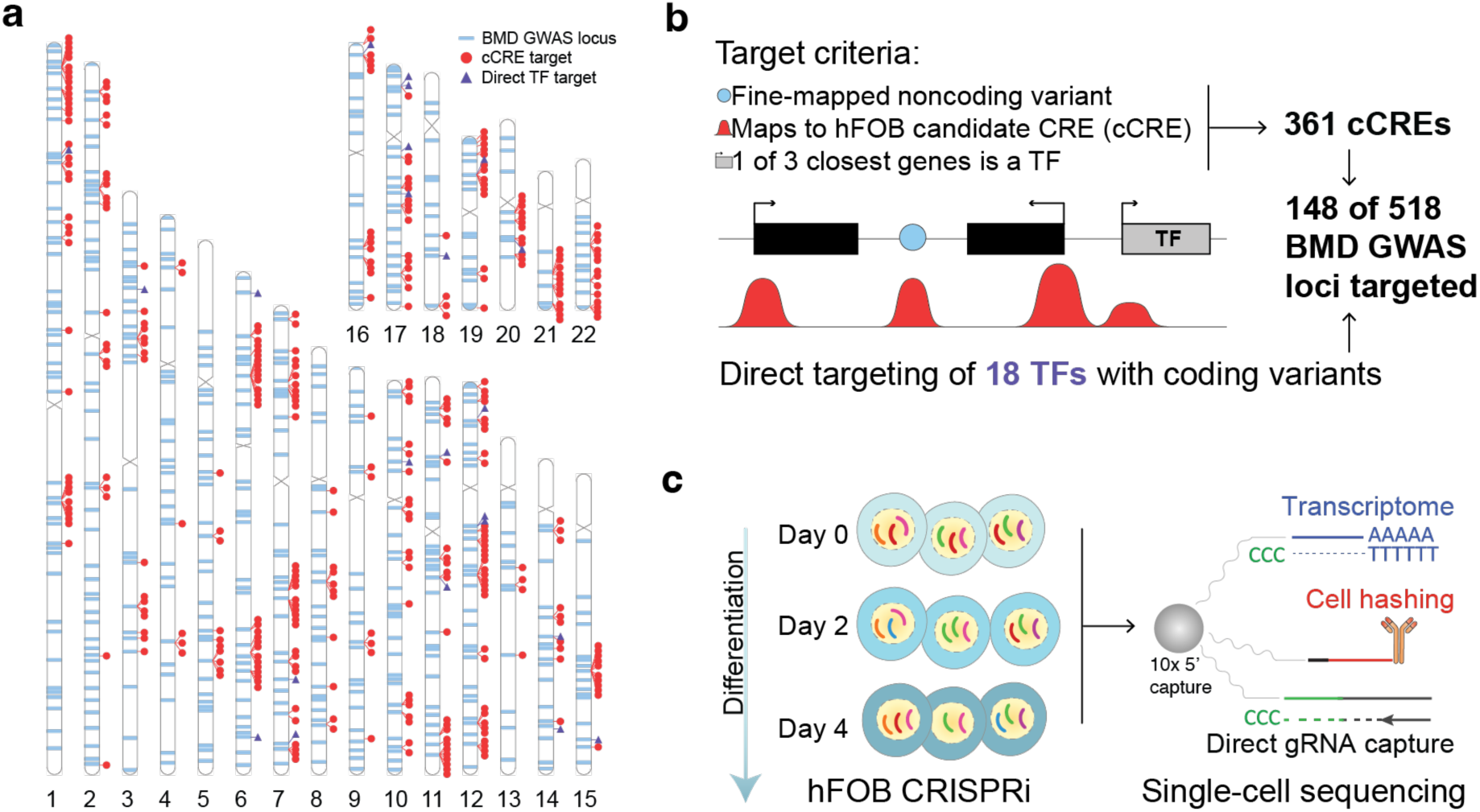
BMD STING-seq guide RNA library (gRNA) and experimental design. **a,** The ideogram shows BMD GWAS loci with targets in the BMD STING-seq gRNA library and labels them if they are cCRE targets in hFOB cells or if they target gene TSSs directly. **b,** The targeting criteria define how the BMD STING-seq gRNA library includes BMD GWAS variants. **c,** We induced hFOB cells to undergo osteogenic induction and differentiation for up to four days after transduction with the BMD STING-seq gRNA library, with multi-modal single-cell sequencing performed at each time point to capture transcriptomes, direct gRNA capture, and cell hashing to improve single-cell data quality.

### BMD STING-seq in hFOB cells

We performed lentiviral transductions of the BMD STING-seq gRNA library in hFOB CRISPRi cells to a high MOI. We selected cells for seven days with puromycin to ensure successful CRISPRi, and then initiated osteogenic differentiation. After an initial pilot experiment on undifferentiated cells, we performed multi-modal single-cell sequencing readouts at days 0, 2, and 4 of osteogenic induction using cell hashing (labeling cells with oligo-tagged antibodies to enable sample multiplexing) to improve single cell recovery and reduce batch effects (**Fig. 1c**) ^46^. Cell hashing enabled multiplexing of multiple time points within the same experiment. Therefore, for each cell, we captured their transcriptomes, gRNAs, and cell hashing antibodies (**Methods**). We performed single-cell sequencing quality control to remove low-quality cells and droplet multiplets (**Methods**), verified a high MOI (median of 8 gRNAs per cell), and assigned gRNAs to individual cells. In total, 77,986 high-quality single cells were retained for downstream analyses: 49,931 undifferentiated day 0 cells (including 23,193 cells from the pilot experiment) and 28,055 cells post-osteogenic induction (17,990 cells from day 2 and 10,065 cells from day 4) (**Extended Data Fig. 4a**, **Supplementary Table 2e**). We performed differential expression testing with SCEPTRE, which was designed for high MOI CRISPR screens ^47^, to identify gRNA–gene pairs and verify successful TSS targeting. We examined five possible time point combinations to discover target genes of cCREs in *cis*, which we define as within 1 Megabase (Mb): days 0, 2, and 4 individually, a combined time point post-induction (days 2 and 4), and all time points combined. For each time point combination, we verified our differentiation expression models were well-calibrated by examining non-targeting gRNAs, finding no spurious associations of non-targeting gRNAs with gene expression (**Extended Data Fig. 4b–f, Supplementary Table 2f**). Next, we examined TSS-targeting gRNAs and found that all 18 genes were significantly inhibited at a 5% false discovery rate (FDR, *q*_SCEPTRE_ < 0.05) by targeting at least one of their TSSs identified from long-read sequencing (**Extended Data Fig. 4g**). TSS-targeting gRNA differential expression tests were also well-calibrated (**Extended Data Fig. 4h–l, Supplementary Table 2g**). Our BMD STING-seq experiment in hFOB cells was therefore well-calibrated across multiple time points. We also inspected individual gRNA effects and found that most TSS-targeting gRNAs exhibited independently significant inhibition of the target genes. For a subset of TSS–gene pairs, we found no single gRNA reached significance on its own, and the association was only detected when combining all three gRNAs to increase power (**Extended Data Fig. 5a**, **Supplementary Table 2h**).

### Target gene identification at noncoding BMD GWAS loci

Across all time points, we identified 91 significant (*q*_SCEPTRE_ < 0.05) CRE–gene pairs in *cis* (**Fig. 2a, Extended Data Fig. 6, Supplementary Table 2i**). In total the CRE–gene pairs identified represented 75 unique target genes. These genes were enriched for gene ontologies such as “skeletal system development” (*q*_FDR_ = 9.1 × 10^-9^) and included several genes with well-established roles in bone biology, such as *DLX5*, *HOXC9*, *PRRX1*, *RUNX1*, and *SMAD3*. The 75 genes were located in 50 (36%) of the 138 noncoding loci targeted. Of the total number of CRE– gene pairs, 64 CREs had a single target gene in *cis* (**Fig. 2b**). Almost 50% of the identified target genes were not the closest gene to their respective CRE (**Fig. 2c**). As expected, CRE–gene distances were greater for target genes that were not the nearest gene (**Fig. 2d**). Nine CREs were within 1 kb of their target gene TSS and were classified as promoter CREs (pCREs). We again inspected individual gRNA effects and found that most CRE–targeting gRNAs exhibited independently significant inhibition of the target genes (**Extended Data Fig. 5b**, **Supplementary Table 2j**).

**Figure 2.**
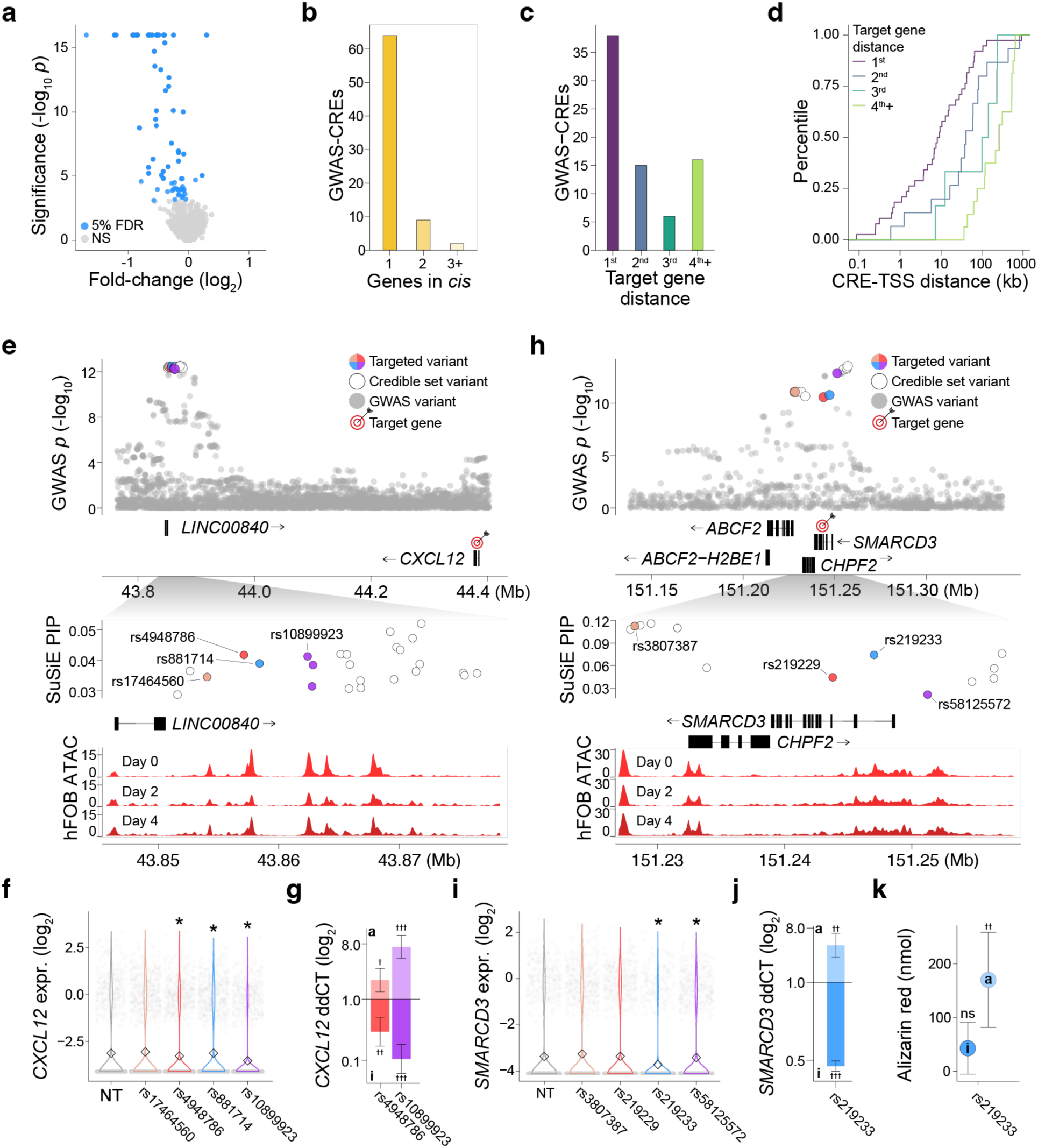
Mapping target genes in cis for noncoding BMD GWAS loci. **a,** The volcano plot displays all CRE–gene pairs across all hFOB time points and identifies 91 significant CRE–gene pairs (5% FDR) up to a distance of 1 Mb. **b,** The number of target genes per CRE. **c,** The distance to gene rank of the closest target gene TSS to its CRE. **d,** The cumulative frequency distribution of the distance between the closest target gene TSS and its CRE. **e,** At this BMD GWAS locus, we targeted four cCREs that we identified from six fine-mapped variants. Variants rs17464560 (salmon) and rs10899923 (purple), along with two other variants, mapped to distinct ATAC-seq peaks, while rs4948786 (red) and rs881714 (blue) mapped to the same ATAC-seq peak but were over 1 kb apart and targeted separately. We identified *CXCL12*, located over 500 kb downstream, as the target gene for this locus. **f,** Single-cell gene expression profiles showed cells where we targeted at rs17464560, rs4948786, rs881714, rs10899923, or non-targeting (NT). Targeted inhibition of the rs4948786, rs881714, and rs10899923 identified CREs significantly decreased *CXCL12* expression. **g,** We validated the rs4948786 and rs10899923 identified CREs for *CXCL12* with CRISPRi and CRISPRa in hFOB cells and measured decreased and increased *CXCL12* expression, respectively. **h,** At this BMD GWAS locus, we targeted four cCREs that we identified from four fine-mapped variants. Variants rs3807387 (salmon), rs219229 (red), rs219233 (blue), and rs58125572 (purple) mapped to distinct ATAC-seq peaks. Two variants lay within introns of *SMARCD3*, which we identified as the target gene for this locus. **i,** Single-cell gene expression profiles showed cells where we targeted at rs3807387, rs219229, rs219233, rs58125572, or NT. Targeted inhibition of the rs219233 and rs58125572 identified CREs significantly decreased *SMARCD3* expression. **j,** We validated the rs219233 identified CRE for *SMARCD3* with CRISPRi and CRISPRa in hFOB cells and measured decreased and increased *SMARCD3* expression, respectively. **k,** We tested for mineralization after targeted inhibition and activation of the rs219233 identified CRE following 8 days of osteogenic induction, followed by Alizarin red staining. Increased *SMARCD3* expression resulted in increased hFOB mineralization. Asterisks denote *q*-values, defined as Benjamini-Hochberg adjusted SCEPTRE *p*-values, significant at a 5% FDR. Daggers denote significant *p*-values from linear mixed models for arrayed experiments, comparing targets against NT († *p* < 0.05, †† *p* < 0.01, ††† *p* < 0.001).

Of the total, 16 (18%) of CRE–gene pairs were time point specific. For example, inhibiting the rs76287541 (C/T, *beta*_GWAS_ = −0.02, *p*_GWAS_ = 8.5 × 10^-14^, PIP_SuSIE_ = 1.0) CRE significantly decreased *NFIC* expression (*p*_SCEPTRE_ = 1.2 × 10^-4^) at day 0, however, the target gene switched, significantly increasing *APBA3* expression (*p*_SCEPTRE_ = 2.0 × 10^-5^), at day 2 (**Supplementary Table 2i**). Inhibiting the rs76287541 CRE had no effect on either gene at day 4 (*p*_SCEPTRE_ > 0.05). *NFIC* encodes a transcription factor that has a binding site at the *APBA3* promoter with evidence from ChIP-seq in ENCODE. However, we mainly report differential expression testing from combining all three time points, as we found these tend to be the most significant due to increased numbers of cells per target (*i.e.*, larger sample size, increased power).

We validated CRE–gene associations across seven selected loci by performing arrayed transductions of the significantly associated gRNAs and, in some cases, newly designed gRNAs targeting the same CREs, using both CRISPRi and CRISPRa in hFOB cells (**Supplementary Table 3a–b**). In these experiments, we measured target gene expression by qPCR and *in vitro* mineralization by calcium deposit staining, a key indicator of osteoblast activity, with Alizarin Red. We confirmed the expression effects observed with STING-seq for six of the seven loci, with CRISPRi decreasing and CRISPRa increasing target gene expression (**Extended Data Fig. 7a, Supplementary Table 3c–d)**. Furthermore, perturbing these CREs altered *in vitro* mineralization at five of the seven loci, establishing connections between fine-mapped variant-tagged CREs, their target genes, and modulation of a bone cell phenotype that directly impacts BMD (**Extended Data Fig. 7b, Supplementary Table 3e–f**). Here, we highlight long-range variant-tagged CREs linked to *CXCL12* (**Fig. 2e–g**) and more proximal variant-tagged CREs linked to *SMARCD3* (**Fig. 2h–k**).

We targeted four cCREs containing fine-mapped variants located just downstream of the long noncoding RNA *LINC00840* (**Fig. 2e**). We found three reduced *CXCL12* expression, located ∼500 kb downstream (**Fig. 2f**). In the bone microenvironment, the chemokine SDF-1 (Stromal Cell-Derived Factor-1; encoded by *CXCL12*) regulates bone mass by retaining CXCR4-positive osteoclast precursors in the marrow niche and promoting osteoblast differentiation; together these actions balance bone resorption and formation and thus influence BMD ^48^. In follow-up experiments, we targeted two CREs, overlapping rs4948786 (G/A, *beta*_GWAS_ = −0.01, *p*_GWAS_ = 4.4 × 10^-13^, PIP_SuSIE_ = 0.04) and rs10899923 (T/C, *beta*_GWAS_ = −0.01, *p*_GWAS_ = 5.2 × 10^-13^, PIP_SuSIE_ = 0.04), with CRISPRi, which again decreased *CXCL12* expression, while CRISPRa increased its expression (**Fig. 2g**). Neither CRISPRi nor CRISPRa altered mineralization (**Extended Data Fig. 7b**), consistent with a role of *CXCL12* in osteoblast–osteoclast coupling rather than direct control of osteoblast differentiation ^49^. Together, these results suggest that long-range CREs modulate *CXCL12* to mediate this BMD association.

We targeted four variant-tagged cCREs in the vicinity of a BMD association near *SMARCD3* (**Fig. 2h**). Targeting two of the four in our STING-seq experiment resulted in the reduced expression of *SMARCD3*. The rs219233 (G/T, *beta*_GWAS_ = −0.01, *p*_GWAS_ = 1.7 × 10^-11^, PIP_SuSIE_ = 0.07) CRE lies within an intron of *SMARCD3*, 1.6 kb downstream of its TSS (**Fig. 2h**). Both CRISPRi and CRISPRa perturbations of the CRE containing rs219233 confirmed bidirectional regulation: CRISPRi reduced and CRISPRa increased *SMARCD3* transcript levels (**Fig. 2i–j**). Importantly, CRISPRa-mediated upregulation of *SMARCD3* significantly enhanced mineralization (**Fig. 2k**), consistent with a positive regulatory role for *SMARCD3* in osteoblast function. Although *SMARCD3* (also known as BAF60C) is a known SWI/SNF chromatin-remodeling subunit essential for cardiac and muscle development, these results identify its first functional connection to osteoblast differentiation ^50^. Together, these data position *SMARCD3* as a strong candidate effector gene underlying BMD association signals at this locus.

### CRISPRi effects replicate in primary hMSCs

To evaluate whether the regulatory effects identified in hFOBs were reproduced in primary human cells, we performed targeted lentiviral CRISPRi in bone-derived hMSCs from a single donor (**Fig. 3a**, **Methods**). Cells were co-transduced with KRAB–dCas9–MeCP2 and individual gRNAs targeting CREs linked to *LINC02454*, *CXCL12*, *SMARCD3*, and *TLN1*, or targeting the *TRIP11* TSS and the *DAP3–YY1AP1* bidirectional promoter (*TRIP11* and *DAP3*–*YY1AP1* discussed below). Target transcript abundance was measured by RT-qPCR in 3–4 independent experiments per guide-transcript pair (**Supplementary Table 3g**). Across 17 matched comparisons, 14 (82%) produced effects in the same direction in hMSCs and hFOBs, and the magnitudes of these effects were strongly correlated (*r*_Pearson_ = 0.83, **Fig. 3b**, **Supplementary Table 3h**). Although effects were generally attenuated in hMSCs, the strong directional and quantitative concordance demonstrates that the regulatory relationships observed in hFOBs were broadly reproduced in primary cells.

**Figure 3.**
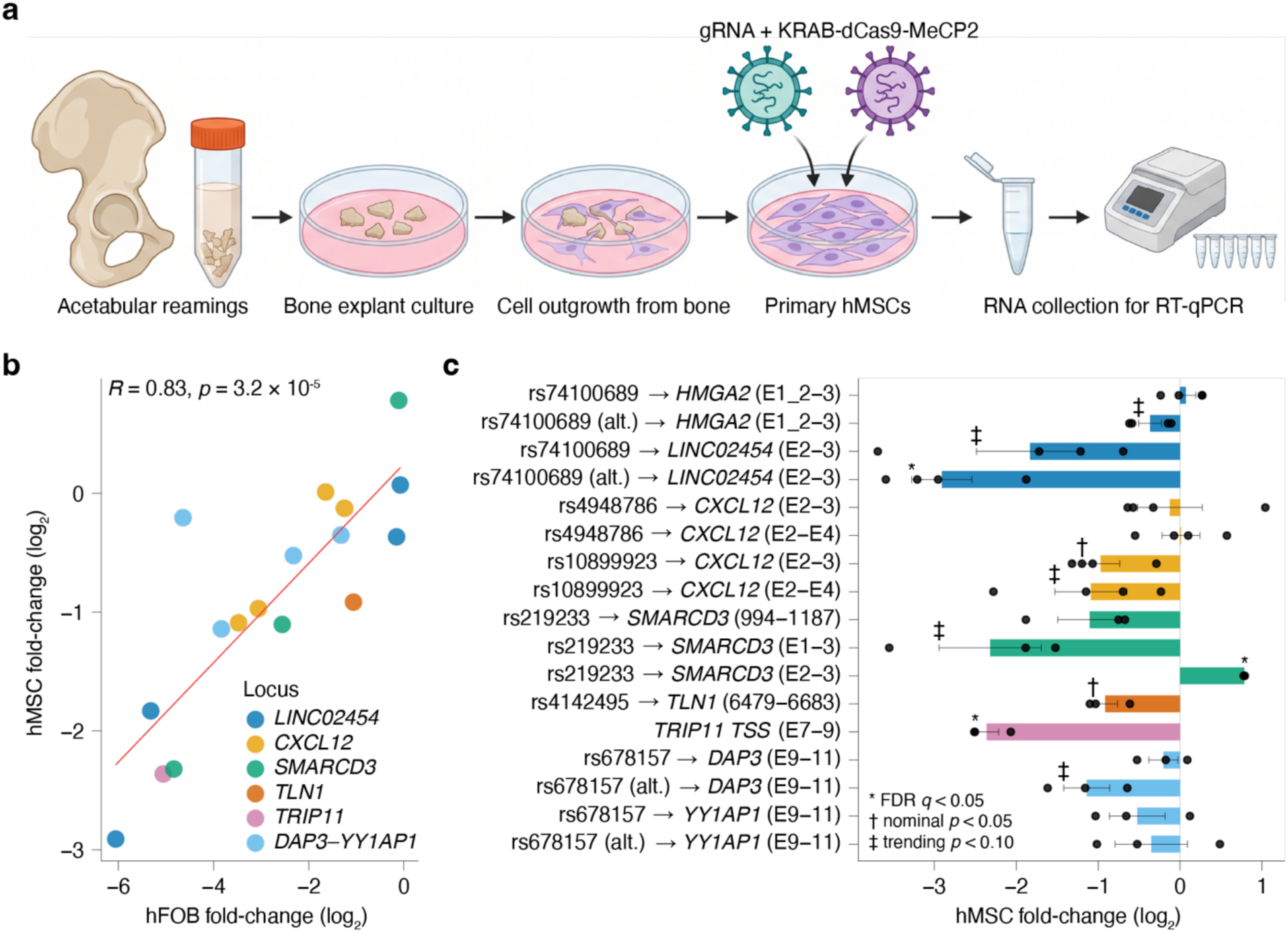
CRISPRi effects are concordant in primary human mesenchymal stem cells (hMSCs). **a,** Primary hMSCs derived from a single donor were co-transduced with lentiviruses encoding a gRNA and KRAB-dCas9-MeCP2 and harvested 3-10 days later for RT-qPCR. **b,** Correlation between CRISPRi effects measured in hFOB cells and primary hMSCs for the 17 gRNA-target comparisons shown in **c**. Each point represents one comparison and is coloured by locus. The red line indicates the linear fit (Pearson *R* = 0.83, *p* = 3.2 × 10⁻⁵). **c,** Effects of CRISPRi on target gene expression in primary hMSCs. Expression was normalized to *CHMP2A* and is shown relative to matched non-targeting controls. Bars show mean log₂ fold-change across three or four independent experiments, bars indicate standard error of the mean, and points indicate individual experiments. Rows correspond to individual gRNAs and assayed transcripts; "(alt.)" denotes an alternative gRNA targeting the same locus. * FDR q < 0.05; † nominal p < 0.05; ‡ p < 0.10. hMSC test statistics were calculated using two-sided t-tests. **a** created in BioRender. Morris, J. (2026) https://BioRender.com/bl86lsi.

Consistent with this overall concordance, several of the largest effects were individually supported in primary hMSCs (**Fig. 3c**). At a 5% FDR, targeting an upstream *LINC02454* CRE reduced *LINC02454* expression by a log_2_ fold change of −2.9 compared to matched non-targeting controls (*q*_FDR_ = 0.024; *n* = 4). This CRE had a secondary effect on *HMGA2* in hFOB cells that we did not observe in hMSCs, but the measured effects were modest in both cases (hFOB log_2_ fold change = −0.01, hMSC log_2_ fold change = 0.01), suggesting a secondary effect on *HMGA2* requires further testing. Direct targeting of the *TRIP11* TSS reduced *TRIP11* expression by a log_2_ fold change of −2.4 (*q*_FDR_ = 0.024; *n* = 3). Targeting CREs linked to *CXCL12* and *TLN1* reduced their expression by log_2_ fold changes of −1.0 and −0.9, respectively (Nominal *p* = 0.025 and 0.027; *q*_FDR_ = 0.091 for both), providing supportive evidence in the expected direction. *SMARCD3* responses were transcript dependent: two amplicons showed decreased expression, whereas the E2–3 amplicon increased (log_2_ fold change = 0.78, *q*_FDR_ = 4.4 × 10^-4^) and was discordant with the effect observed in hFOB cells (hFOB log_2_ fold change = −0.1). Together, these results show that the pattern and relative magnitude of CRISPRi effects are reproducible between hFOBs and primary hMSCs, while also identifying locus- and transcript-specific effects that should be resolved in additional donors.

### Linking rs67631072 to *FHL3* with STING-seq and allele specific open chromatin

Given that we identified significant CRE–gene pairs from fine-mapped noncoding GWAS variants, we next sought to pinpoint variants that may be directly functional in hFOBs using allele-specific regulatory analyses. Whole-genome sequencing of hFOBs identified 189 heterozygous variants out of the 570 used to design the BMD STING-seq gRNA library. Inspection of allele-specific ATAC-seq read counts across all time points (**Extended Data Fig. 8a**) revealed rs67631072 (C/T, *beta*_GWAS_ = −0.02, *p*_GWAS_ = 7.7 × 10^-25^, PIP_SuSIE_ = 0.2), which we found had consistent allelic imbalance across each time point (**Extended Data Fig. 8b**). STING-seq linked this variant-tagged CRE to *FHL3*, which showed a modest but statistically significant expression change (*p*_SCEPTRE_ = 3.1 × 10⁻⁴) (**Extended Data Fig. 8c**). The variant lies within a CRE located ∼600 bp downstream of the *FHL3* 3′ UTR and ∼9 kb from its TSS (**Extended Data Fig. 8d**). Although previously annotated to *SF3A3*, the nearest gene, and itself a regulator of *RUNX2* ^51^, our functional genomic data instead nominates *FHL3* as the target gene, with no significant change in *SF3A3* expression (*p*_SCEPTRE_ = 0.4). We previously used splicing quantitative trait loci (sQTLs) ^52^ and expression QTLs (eQTLs) ^53^ from 49 GTEx tissues and performed genome-wide colocalization for GTEx sQTLs and eQTLs with our BMD GWAS ^31^. This analysis revealed *FHL3* as one of the strongest sQTL and eQTL colocalizations with BMD, and with rs67631072 as the lead variant. Here, we report an updated colocalization between our targeted variants and eQTLs from GTEx, again colocalizing *FHL3* with BMD, and with rs67631072 also as the lead eQTL variant (**Supplementary Table 4**). We further inspected the rs67631072 allele change (C-to-T) and found it disrupted an NF-κB binding site, which also has a known role in bone formation ^54^. Together, the allele-specific chromatin imbalance at rs67631072, the CRE–gene assignment from STING-seq, and sQTL colocalization converge on *FHL3* as the target gene mediated by rs67631072 at this noncoding BMD locus (**Extended Data Fig. 8d**).

### *Trans*-regulatory networks reveal insight into the function of *cis* target genes

To investigate how target genes in *cis* potentially influence BMD through regulatory networks, we searched for genes acting downstream of the 75 identified *cis*-target genes (**Fig. 4a**). We identified *trans*-regulatory networks that influenced the expression of at least 10 downstream targets for five target genes or gene pairs: *TRIP11*, *DAP3*–*YY1AP1*, *UBP1*, *ZBTB4*, and *CHD4* (**Fig. 4b, Extended Data Fig. 6, Supplementary Table 5a**). We examined individual gRNA effects and found their effects on downstream target genes were positively correlated across all comparisons, indicating that *trans*-regulated networks are not driven by gRNA specific effects or seed region off-targets (**Extended Data Fig. 9**). All five *trans*-regulatory networks showed enrichment for multiple biological functions, including strong enrichment for the *TRIP11* network in Golgi apparatus function (*p*_Fisher’s_ = 1.0 × 10^-22^) and the *DAP3*–*YY1AP1* network in oxidative phosphorylation (*p*_Fisher’s_ = 6.0 × 10^-12^) (**Supplementary Table 5b**) ^55^. These enrichments provide insight into how these genes may influence osteoblast activity and BMD.

**Figure 4.**
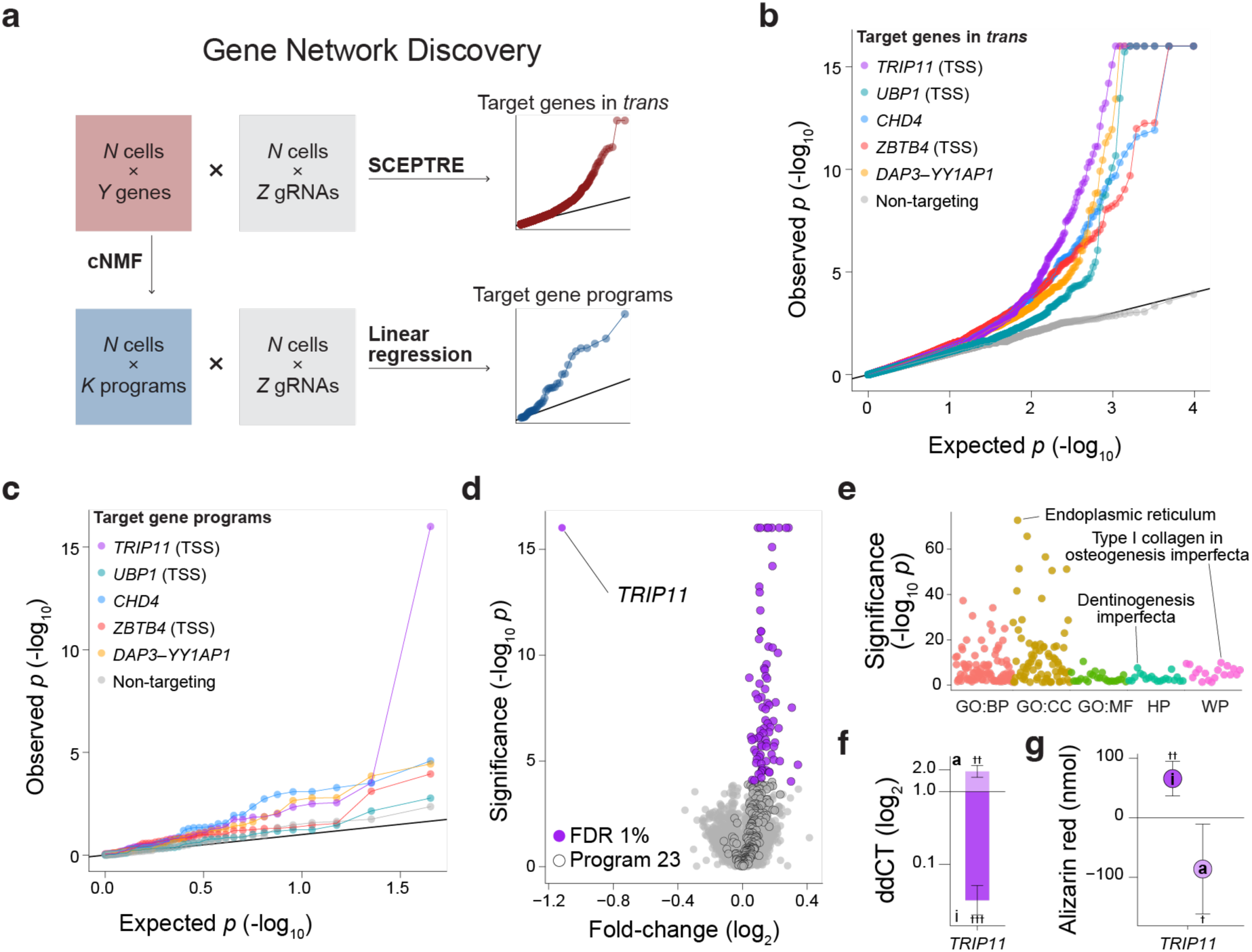
Gene network discovery for targeted CREs. **a,** We used two complementary methods for gene network discovery: transcriptome-wide differential gene expression testing and cNMF gene program testing. **b,** The QQ-plot shows targeted CREs or TSSs that had multiple significant target genes (1% FDR), defined as at least 10 target genes in *trans.* **c,** The QQ-plot shows targeted CREs or TSSs that had a significant association with cNMF programs (1% FDR). Programs were constructed with cNMF in sets of 300 genes. **d,** Volcano plot of the *TRIP11* gene network. Including *TRIP11*, we identified 95 significant differentially expressed genes (1% FDR, purple). Eighty-one of the 95 genes were members of cNMF program 23, which was also significantly associated with *TRIP11* TSS targeting (1% FDR, black circle). **e,** Gene set enrichments for *TRIP11* network genes highlighted roles in Golgi apparatus function and bone-related functions and disorders. **f,** We performed arrayed CRISPRi and CRISPRa targeting of the *TRIP11* TSS to validate changes in expression. **g,** We quantified changes in hFOB mineralization and found that *TRIP11* inhibition increased mineralization, whereas *TRIP11* activation decreased mineralization. Daggers denote significant *p*-values from linear mixed models for arrayed experiments, comparing targets against NT († *p* < 0.05, †† *p* < 0.01, ††† *p* < 0.001).

To generate a more fine-scaled view of transcriptional regulation, we used consensus non-negative matrix factorization (cNMF) to identify gene programs ^22,56,57^. cNMF decomposes single-cell expression profiles into sets of genes that are co-expressed across cells. Each gene program represents a shared transcriptional state or biological process, and each cell receives a program usage score that reflects the extent to which that program is active in the cell. cNMF has been shown to recover biologically meaningful programs associated with metabolic functions, developmental states, and cell identity ^22,56,57^. We applied cNMF and identified 45 hFOB gene programs (**Supplementary Table 5c**), then tested all gRNAs for associations with usage of transcriptional programs (**Fig. 4a**). We confirmed our analyses were well-calibrated (**Supplementary Table 5d**) and identified 33 gRNA–program associations at a 1% FDR (**Supplementary Table 5e**). We focused on a high-confidence subset of gRNA–program associations—those where the gRNAs also had significant *trans*-regulatory networks—and identified 11 high-confidence associations: *TRIP11* with programs 23 and 30, *CHD4* with programs 2, 20, 34, 35, 43, and 44, *DAP3*–*YY1AP1* with programs 20 and 31, and *ZBTB4* with program 30 (**Fig. 4c**). For example, based on program associations, we can infer that *CHD4*, a core ATP-dependent chromatin remodeler, may impact osteoblast activity by altering multiple coordinated transcriptional programs. We linked perturbation of *CHD4* to programs associated with cell cycle progression (Programs 2 and 34), matrix and secretory pathways (Program 35), and transcriptional regulatory modules (Programs 20 and 44), as well as signatures related to ribosomal and housekeeping functions (Program 43) (**Supplementary Table 5e–f**). These associations suggest that *CHD4* influences chromatin accessibility across diverse functional axes relevant to osteoblast biology, integrating regulation of proliferation, extracellular matrix dynamics, and gene regulatory networks that are critical for osteoblast differentiation and function. Together, these results demonstrate orthogonal methods, in this case SCEPTRE and cNMF, can consistently identify genes that have regulatory effects in *trans*. We use these results to further demonstrate below that STING-seq can identify biologically meaningful regulatory networks downstream of BMD GWAS associations.

### *TRIP11* regulates a tunable protein secretion network that regulates osteoblast activity

In our STING-seq screen, we directly targeted the *TRIP11* TSS as it was identified via a non-synonymous fine-mapped BMD variant, rs1051340 (p.Gly1827Cys, C/T, *beta*_GWAS_ = −0.01, *p*_GWAS_ = 6.5 × 10^-9^, PIP_SuSIE_ = 0.02). We observed that direct inhibition of *TRIP11* regulated the expression of 94 other genes as part of a downstream *trans*-regulatory gene network (**Fig. 4d**, **Supplementary Table 5a**) significantly enriched for genes involved in protein transport and secretion (*p*_Fisher’s_ = 1.3 × 10^-14^) (**Fig. 4e**, **Supplementary Table 5b**). These observations were consistent with *TRIP11* encoding the Golgi-microtubule-associated protein GMAP-210. As described above the *TRIP11* perturbation was associated with cNMF program 23. There was a large overlap between the *TRIP11* network and program 23 (81/95 genes found in program 23). In program 23, decreased *TRIP11* expression corresponded with increased expression of most genes in the program (272/300 genes, 81/81 significant) (**Fig. 4d**). This program also contained several key extracellular matrix proteins, including type I collagen (COL1A1), and it showed strong enrichment for Mendelian disease genes that cause osteogenesis imperfecta (**Fig. 4e**, **Supplementary Table 5b**).

We replicated the *TRIP11* gene network in hMSCs to demonstrate that the network is not an hFOB-specific artifact. As discussed above, we demonstrated that we can inhibit *TRIP11* expression in hMSCs using lentivirally delivered CRISPRi and *TRIP11* TSS-targeting gRNAs. We then performed bulk RNA-seq of hMSCs at days 6 and 10 post-lentiviral transduction to capture changes in the *TRIP11* regulatory network (**Methods**). We compared TSS-targeting cells against hMSCs transduced with non-targeting gRNAs and performed our experiments in triplicate. We examined the 95 genes (including *TRIP11*), tested for concordant log_2_ fold-changes between hMSC and hFOB experiments, and found 87% and 88% had log_2_ fold-changes in concordant directions at days 6 and 10, respectively (**Supplementary Table 5g**). We repeated the experiment with *TRIP11*-targeting lentiviral shRNAs compared to hMSCs transduced with empty vectors and again found 84% and 63% had log_2_ fold-changes concordant at days 6 and 10, respectively (**Supplementary Table 5h**). This supports that the observed *TRIP11* network is not an hFOB-specific artifact and is conserved in primary cells.

These findings suggested a functional role for *TRIP11* in osteoblast activity. Targeting the *TRIP11* TSS confirmed that CRISPRi reduced *TRIP11* expression, while CRISPRa increased *TRIP11* expression (**Fig. 4f**). Consistent with the increase in extracellular matrix gene expression observed in program 23, reduced *TRIP11* expression increased mineralization, whereas increased *TRIP11* expression reduced mineralization (**Fig. 4g**). These data suggest that *TRIP11* in osteoblasts regulates a tunable *trans*-regulatory network that modulates mineralization and, possibly, BMD.

### A promoter CRE for *DAP3* and *YY1AP1* regulates a mitochondrial *trans*-regulatory network and mineralization through *DAP3*

We identified a secondary association signal at the previously identified *THBS3* locus, where non-genome-wide significant variants were fine-mapped as plausibly causal. Conditional association testing revealed that rs670876 (T/G, *beta*_GWAS_ = −0.005), the lead fine-mapped variant in the secondary signal (PIP_SuSIE_ = 0.04), became more significant upon conditioning on the top variant (rs914615, A/G, *beta*_GWAS_ = 0.01, *p*_GWAS_ = 4.0 × 10^-13^, PIP_SuSIE_ = 0.04), with *p*_GWAS_ = 1.3 × 10^-2^ and *p*_COJO_ = 1.7 × 10^-4^, providing supporting evidence for this secondary signal. We targeted a cCRE harboring rs678157 (T/C, *beta*_GWAS_ = −0.005, *p*_GWAS_ = 2.1 × 10^-3^, PIP_SuSIE_ = 0.01), which is in near-perfect linkage disequilibrium with rs670876 (r^2^ = 0.98). This CRE functions as a promoter CRE (pCRE) within a bidirectional promoter regulating both *DAP3* and *YY1AP1* (**Fig. 5a**), and its targeting significantly (5% FDR) inhibited expression of both genes (**Fig. 5b**). We also observed that upon arrayed transduction with CRISPRi, there was a near complete loss of mineralization (**Fig. 5c–d**). YY1AP1 is a cofactor for the YY1 transcription factor, which coordinates the regulation of genes involved in growth, differentiation, and apoptosis ^58^. YY1 has also been shown to regulate osteoblast differentiation through interactions with p38 and RUNX2 ^59^. DAP3, a component of the 28S small subunit of the mitochondrial ribosome, is essential for mitochondrial protein synthesis and has been shown to influence mitochondrial morphology when knocked down in HeLa cells ^60^.

**Figure 5.**
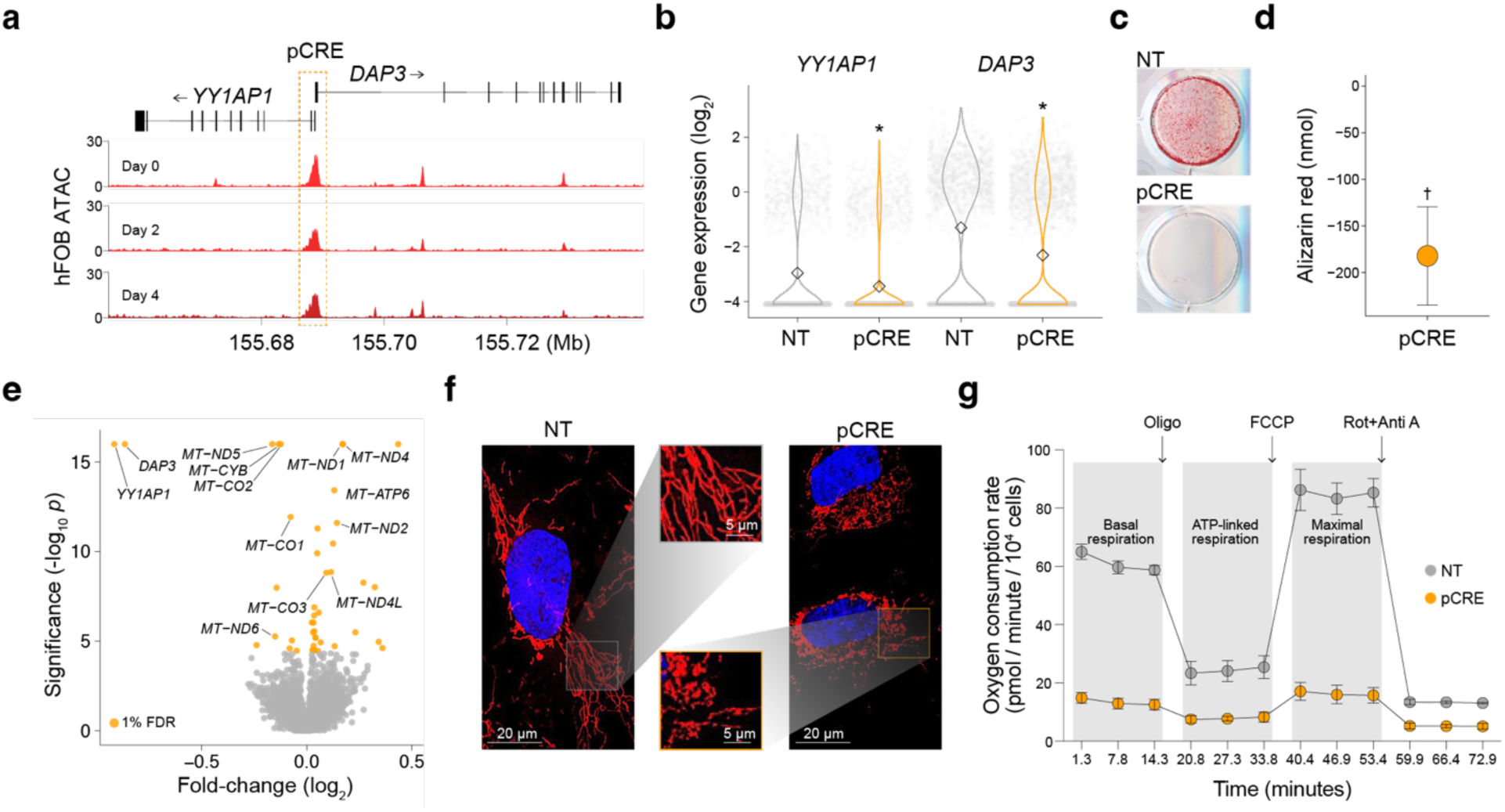
Targeting the *DAP3*–*YY1AP1* locus resulted in loss of mineralization mediated by mitochondrial function. **a,** We targeted a promoter CRE (pCRE) that mapped between the promoters of *DAP3* and *YY1AP1*. **b,** Single-cell gene expression profiles showed cells with gRNAs targeting the pCRE had significantly reduced *YY1AP1* and *DAP3* expression (5% FDR, asterisk), whereas non-targeting (NT) gRNAs had no effect. Targeting the pCRE with arrayed CRISPRi resulted in a complete loss of hFOB mineralization **(c, d)**. **e,** Volcano plot of the *DAP3*– *YY1AP1* gene network. Excluding *DAP3* and *YY1AP1*, we identified 41 significant differentially expressed genes (1% FDR, gold). Eleven of 13 known mitochondrial (MT) genes were identified as part of this gene network. **f,** We performed mitochondrial staining and imaging on arrayed CRISPRi pCRE and NT hFOB cells and observed increased mitochondrial fragmentation upon *DAP3* and *YY1AP1* inhibition. **g,** We measured the cellular oxygen consumption rate over time in arrayed CRISPRi pCRE and NT hFOB cells and observed a substantially reduced ability to consume oxygen across respiration inflection points, including basal, ATP-linked, and maximal respiration, following stimulation with oligomycin (ATP synthase inhibitor), FCCP (mitochondrial uncoupler), and rotenone plus antimycin (Complex I and III inhibitors), respectively. Asterisks denote *q*-values, defined as Benjamini-Hochberg adjusted SCEPTRE *p*-values, significant at a 5% FDR. Daggers denote significant *p*-values from linear mixed models for arrayed experiments, comparing targets against NT († *p* < 0.05).

As described above, we identified a *trans*-regulatory network of 41 differentially expressed genes at a 1% FDR (**Fig. 5e**). This network contained nearly all (11 of 13) mitochondrially encoded members of the electron transport chain. Given that mineralization is an energy-intensive process, this supports our finding that the *DAP3*–*YY1AP1* pCRE is essential for mineralization. Continuing this line of evidence, and the involvement of *DAP3* in mitochondrial function ^60^, we assessed mitochondrial morphology and function in hFOB cells where we targeted the pCRE. Imaging revealed increased mitochondrial fragmentation following pCRE inhibition (**Fig. 5f**). Seahorse assays, that measure oxygen consumption rate, further demonstrated that these changes were accompanied by a significant reduction in oxidative capacity in cells with decreased *DAP3*– *YY1AP1* expression (**Fig. 5g**). We also designed an additional gRNA, not taken from the BMD STING-seq library, to target the pCRE, replicating our findings with this independent gRNA in basal media and osteogenic media (**Extended Data Fig. 10a–d, Supplementary Table 6a–h**).

Next, we performed arrayed siRNA transfections to independently knock down *DAP3* and *YY1AP1* and assess their individual effects on mineralization and mitochondrial function in hFOB cells. Using this approach, we successfully downregulated *DAP3*, whereas *YY1AP1* knockdown was not successful (**Extended Data Fig. 10e–g, Supplementary Table 6i–k**). Repeating mineralization assays following *DAP3* siRNA knockdown fully recapitulated the effects observed with targeting of the *DAP3*–*YY1AP1* pCRE (**Extended Data Fig. 10h, Supplementary Table 6l– m**). Thus, while *YY1AP1* may have independent roles in osteoblast activity and BMD, the mitochondrial-mediated effects observed here are directly driven by *DAP3*. We next tested whether the *DAP3* network could be replicated in primary hMSCs using lentiviral shRNAs. Consistent with the hFOB results, the majority of network genes showed concordant directions of effect in hMSCs (76.2% and 73.8% concordant at days 6 and 10, respectively; **Supplementary Table 5i**), supporting that the observed network is not specific to hFOB cells. However, shRNA-mediated repression of *DAP3* in hMSCs was also accompanied by reduced *YY1AP1* expression, preventing definitive separation of their individual contributions in hMSCs. Together, these data link a secondary BMD GWAS signal to the expression of *DAP3*, which in turn regulates a *trans*-regulatory network that controls mitochondrial and osteoblast activity.

## Discussion

The most pressing challenge in complex disease genetics is translating genetic associations into biological function by connecting disease-associated variants to the genes and regulatory pathways they affect (V2F). Overcoming the V2F gap is essential for enabling rational therapeutic discovery and precision medicine, yet remains particularly difficult for complex traits, such as BMD, where the vast majority of associated variants reside in noncoding regulatory regions. In this study, we address this challenge by applying STING-seq to systematically link fine-mapped BMD variants from large human cohorts to their target genes through CREs, while simultaneously revealing the downstream *trans*-regulatory networks they control in a disease-relevant osteoblast model. By applying STING-seq across osteogenic differentiation, our approach connects BMD-associated variants in human populations to their target genes and downstream biological functions at scale, providing new insight into both the local regulatory effects of individual variants and their broader influence on osteoblast gene regulatory networks.

Molecular quantitative trait locus identification (*e.g.*, eQTLs, sQTLs) and colocalization have proven powerful approaches for identifying putative target genes downstream of GWAS loci ^61^. However, such studies remain scarce in bone and bone-relevant cell types, limiting the ability of colocalization approaches to identify target genes. Consequently, prior efforts to map BMD-associated variants to target genes have largely relied on GTEx eQTL and sQTL resources, which are derived from bulk adult tissues and may not capture regulatory programs active in osteoblasts or their progenitors ^31,53^. In this context of limited disease-relevant regulatory maps, our application of STING-seq is particularly valuable. Using STING-seq, we linked variant-tagged CREs to 75 putative target genes, many of which have not previously been implicated in bone biology and therefore represent high-priority candidates for future functional studies. Most CREs were linked to a single target gene, although a subset mapped to two or more genes. Importantly, CRISPRi of a variant-tagged CRE does not reproduce the effect of an individual GWAS allele. GWAS effect estimates measure allelic associations with BMD, whereas CRISPRi inhibits an entire regulatory element and may produce substantially larger effects; these effects are therefore not directly comparable in magnitude or direction. Of the 361 cCREs targeted in this study, 16 contained more than one fine-mapped variant, including seven for which we identified at least one significant target gene. Although our experiments do not distinguish which variant within a cCRE is causal or whether multiple variants contribute to its activity, they establish functional regulatory connections between variant-tagged CREs and their target genes, substantially narrowing the candidates for future allele-specific studies.

We observed that for more than 45% of CREs linked to a target gene, the inferred target was not the closest gene to the lead variant. This proportion is substantially higher than the approximately 13% (18 of 134) reported in our prior STING-seq study of blood traits ^6^. This difference may reflect a distinct feature of the genetic architecture of BMD, in which long-range regulatory interactions play a prominent role, or it may arise from methodological differences, including the chromatin features used to prioritize regulatory elements for interrogation (*e.g.*, ATAC-seq alone versus combinations of ATAC-seq and H3K27ac ChIP-seq). Consistent with this interpretation, many of the CREs we targeted function as long-range enhancers, with 24 CRE–TSS distances exceeding 100 kb and 9 exceeding 500 kb, including the *CXCL12* locus (**Fig. 2e**). Together, these findings underscore the value of STING-seq and provide a curated set of candidate effector genes for future mechanistic and *in vivo* studies of skeletal biology.

A central challenge in V2F studies is understanding how causal genes influence downstream *trans*-regulatory networks to alter cellular function and ultimately contribute to disease risk. This problem is particularly acute for complex traits, such as BMD, where thousands of disease-associated variants likely act in aggregate by perturbing discrete components of cellular regulatory networks rather than single genes or pathways. Defining these downstream networks is therefore essential both for mechanistic insight and for identifying therapeutic strategies that selectively target disease-relevant biological processes.

STING-seq-based perturbations provided a powerful framework to interrogate such networks in a systematic manner. Building on our prior work ^6^, we focused on disease-associated loci containing TFs or regulatory elements predicted to control transcriptional programs in *trans*, with the goal of identifying the downstream networks regulated by these loci. Using this approach, we identified *trans*-regulatory networks for five genes, most of which were perturbed by targeting either the gene TSS or a pCRE. As an orthogonal and complementary strategy, we applied cNMF to identify gene expression programs and then associated gRNAs with these programs. This analysis provided a more granular view of the regulatory architecture downstream of each target gene, enabling us to connect individual perturbations to coherent biological processes and to infer potential gene functions from the transcriptional programs they regulate. STING-seq enabled us to connect disease-associated variants to CREs, from regulatory elements to target genes, and from target genes to the downstream networks they control; thereby completing the V2F linkage. Importantly, we were also able to directly validate inferred functions for two of the identified targets, providing functional confirmation of these regulatory connections.

For *TRIP11*, integration of *trans*-regulatory networks and cNMF program associations suggested a role in regulating protein secretion in osteoblasts, consistent with its known function as a Golgi-associated tethering protein ^62^. Extending this inference, we demonstrated that perturbation of *TRIP11* directly altered osteoblast mineralization. Specifically, reducing *TRIP11* expression led to increased mineralization, whereas *TRIP11* overexpression suppressed mineralized nodule formation. In chondrocytes, loss-of-function mutations in *TRIP11* cause selective intracellular retention of a subset of cartilage-specific extracellular matrix proteins, while other secreted proteins traffic normally ^63^. Together, these observations suggest that *TRIP11* functions as a negative regulator of osteoblast mineralization, potentially through selective retention of proteins that play critical roles in matrix maturation or mineral deposition. Further studies will be required to define the precise cargo and trafficking mechanisms underlying this effect.

We also identified a pCRE whose perturbation simultaneously affected *DAP3* and *YY1AP1*, resulting in disruption of nearly all genes (11 of 13) encoding components of the electron transport chain. Consistent with this transcriptional signature, perturbation of this regulatory element impaired osteoblast mineralization, altered mitochondrial morphology, and reduced mitochondrial respiration. Interpretation of this locus is complicated by the fact that the targeted pCRE overlaps a bidirectional promoter and therefore has the potential to influence the expression of both genes. Nevertheless, the observed phenotype is mechanistically plausible, as *DAP3* is a core component of the mitochondrial ribosome. Using targeted perturbations, we further interrogated the contribution of each gene independently and found that *DAP3* knockdown recapitulated the major effects on mitochondrial function and mineralization observed following pCRE perturbation in hFOB cells. Together, these findings implicate *DAP3* as the primary driver of the phenotype downstream of the shared regulatory element, while not excluding a contribution from *YY1AP1*.

We recognize that as our discovery screens and initial validation experiments were performed in hFOB cells, it remains possible that some results reflect hFOB-specific behavior rather than true osteoblast differentiation states. We have attempted to mitigate this possibility by replicating CRE–gene pairs in primary hMSCs, with an 82% success rate, and observed that the *TRIP11* and *DAP3–YY1AP1 trans*-regulatory networks are conserved in hMSCs. We have also inspected our *trans*-regulatory network results for possible CRISPRi-specific off-target effects, based on recent findings that NGG motifs in the 12-nucleotide seed region of a gRNA sequence results in promiscuous inhibitory effects ^40–42^. We found this was unlikely in our case, as individual gRNAs, with different sequences, targeting the same CRE or TSS resulted in the same *trans*-regulatory networks (**Extended Data Fig. 9**). Another important caveat of our study is that BMD is influenced by multiple skeletal cell types. While BMD-associated loci are most strongly enriched in osteoblast lineage cells, recent single-cell studies have also demonstrated significant enrichment in chondrocytes, endothelial cells, and vascular smooth muscle cells ^26^. Our usage of ATAC-seq data from hFOB cells may bias CRE nomination towards osteoblast-active regulatory elements and reduce sensitivity for identifying causal regulatory mechanisms in other BMD-relevant cell types. Furthermore, the chromatin landscape of a single osteoblast-like cell line may not fully reflect the regulatory architecture of healthy or osteoporotic bone *in vivo*. Finally, because the loci interrogated in this study were selected from GWAS of estimated BMD derived from heel quantitative ultrasound measurements in predominantly Northern European populations ^2^, our analyses capture only a subset of the genetic and regulatory architecture underlying skeletal traits. Additional loci, regulatory mechanisms, and target genes may emerge from studies of other bone phenotypes and increasingly diverse populations. Nevertheless, this work provides a framework for systematically dissecting the regulatory mechanisms linking BMD GWAS loci to target genes and downstream cellular programs across the broader cellular and genetic landscape of bone biology.

In summary, we utilized STING-seq to systematically link BMD-associated genetic variation to target genes and regulatory networks that govern osteoblast function. This analysis connected fine-mapped noncoding variants to their CREs and target genes and further revealed downstream *trans*-regulatory programs that influence key osteoblast processes, including protein secretion, mitochondrial function, and mineralization. Through functional validation, we identified previously unrecognized roles for genes such as *SMARCD3*, *TRIP11*, and *DAP3* in human bone biology, alongside long-range regulatory control of established BMD genes such as *CXCL12*. Together, these findings provide mechanistic insight into how noncoding GWAS variants shape osteoblast activity and highlight specific genes and pathways mediating genetic effects on BMD.

## Data availability

hFOB ATAC-seq sequencing reads and called peaks at time points 0, 2, 4, and 10 [*day 10 PENDING*] are available at GSE318537. hFOB BMD STING-seq single-cell sequencing reads and UMI count matrices (cDNA, gRNA, and hashtags), all time points combined, are available at GSE319266. hFOB whole-genome sequencing is available through dbGaP at phs004654.v1.p1. Previously generated hFOB short-read and long-read RNA-seq data at time points 0, 2, 4, and 10 at GSE186922 and GSE224588, respectively. hMSC short-read RNA-seq data is available at [*PENDING*].

## Code availability

Custom code used to design the gRNA library, process the single-cell sequencing data, run the SCEPTRE differential expression analyses, and generate figures in this study is available at https://github.com/morrislab-ccbr. Code for the remaining analyses and figures is available from the corresponding authors upon reasonable request.

## Acknowledgements

We thank the entire Farber and Morris laboratories for support and advice. We thank Dr. Timothy Barry for his helpful discussions with implementing cNMF analyses. We thank Dr. Timothy Bullock at the University of Virginia for access to the Seahorse XFe96 analyzer used in this study. This research has been conducted using the UK Biobank Resource under Application Number 47976. Research reported in this publication was supported in part by the National Institute of Arthritis and Musculoskeletal and Skin Diseases of the National Institutes of Health (NIH) under Award Numbers R01AR071657 and R01AR077992 to C.R.F, the NIH under Award Number T32GM145443 to L.W., and by SIF195 Grand Challenge Research Investment: Precision Health for Populations Initiative to C.R.F. J.A.M. was supported by the University of Toronto (U of T), the U of T Connaught Fund, and research infrastructure funded by the Canada Foundation for Innovation (CFI) John R. Evans Leaders Fund and the Ontario Research Fund Small Infrastructure Fund (ORF SIF) under Project Number 46612. H.J.P. was supported by the Canadian Institutes of Health Research CGRS-M Program. C.B. was supported by the U of T Summer Undergraduate Data Sciences Research Opportunity Program. N.E.S. was supported by the National Heart, Lung, and Blood Institute and the National Human Genome Research Institute under Award Numbers R01HL168247 and R01HG012790. The content is solely the responsibility of the authors and does not necessarily represent the official views of the National Institutes of Health or other funders. The funders had no role in study design, data collection and analysis, decision to publish or preparation of the manuscript.

## Contributions

J.A.M. and C.R.F. conceived, designed, and supervised the study. L.M. and J.A.M. designed and performed the single-cell CRISPR screening. Me.K. contributed to CRISPR library off- and on-target scoring analysis. Mo.K. and J.A.M. analyzed single-cell CRISPR screen data and performed downstream analyses. V.M. performed mitochondrial phenotyping assays and related analyses. C.B. performed analyses related to bulk RNA-seq. L.M., L.W., K.K., H.J.P., M.G., A.C.M. and S.G. performed experiments and analyses related to ATAC-seq, allele-specific variant effects, and variant fine-mapping. L.M. and V.M. performed primary hMSC experiments. T.L. and N.E.S. contributed to study design. J.A.M. and C.R.F. wrote the manuscript. All authors read and approved the final manuscript.

## Methods

### Human fetal osteoblast (hFOB) growth conditions

We cultured hFOB 1.19 cells (ATCC CRL-11372) at 34 °C and differentiated them at 39.5 °C as recommended, with the following modifications. For growth media, we used Minimal Essential Media (MEM, Gibco 10370-021) supplemented with 10% Fetal Bovine Serum (FBS, Atlantic Biological S12450), 1% Glutamax (Gibco 35050-061), and 1% Pen Strep (Gibco 15140-122). We seeded cells at a ratio of 1:4–1:6 and passaged them or changed the media every 3 days to avoid extended periods of confluency, which reduce mineralization potential. For differentiation media, we used MEM alpha (Gibco 12571-063) supplemented with 10% FBS, 1% Glutamax, 1% Pen Strep, 50 µg/µL Ascorbic Acid (Sigma A4544-25G), 10 mM beta-Glycerophosphate (Sigma G9422-100G), and 10 nM Dexamethasone (Sigma D4902-25MG).

### hFOB differentiation

Forty-eight hours after plating cells at a density of 75,000 cells per well in a 24-well plate, we removed the growth media and replaced it with 0.5 mL of pre-warmed differentiation media. We then placed the cells at 39.5 °C. We replaced the differentiation media every other day until we determined mineralization deposition (days 7–10). On that day, we triple-washed the cells with 1 mL DPBS per well, fixed them with 0.5 mL of 10% Buffered Formalin Phosphate (Fisher #SF100-4) for 15 minutes at room temperature (RT), and triple-washed them again with 1 mL per well of water. We then stained the cells with 400 µL per well of 40 mM Alizarin Red, pH 5.6 (adjusted with NH₄OH; Sigma A5533-25G) for 25 minutes at RT with gentle shaking. After removing the staining solution, we washed the cells 10 times for 10 minutes each with 1 mL per well of water and gentle shaking at RT. After scanning the plates to collect images, we quantified the amount of Alizarin Red bound to the mineral by eluting it in 2 mL per well of 5% (v/v) Perchloric Acid (HClO₄, Sigma-Aldrich 311413-500ML) and incubating for 20 minutes with gentle shaking at RT. We determined the amount of Alizarin Red bound for each sample or treatment from the absorbance at 405 nm of the eluent, using standards prepared from the Alizarin Red staining solution diluted into 5% HClO₄ (blanked against 5% HClO₄ using a BioRad SmartSpec Plus spectrophotometer), and expressed the results in units of nmol of Alizarin Red bound.

### Detaching viable hFOB cells

We detached cells used for single-cell RNA-seq and ATAC-seq from the plates they were grown in using the following protocol (described for a 6-well culture dish). We removed the growth media, washed the cells with 1 mL Hank’s Balanced Salt Solution (HBSS; Gibco 14025-092), and incubated them with 0.5 mL of 8 mg/mL collagenase (Gibco 17018-029) in HBSS supplemented with 3 mM CaCl₂ (final [Ca²⁺] = 4 mM) for 10 minutes at 37 °C with gentle shaking. We then triturated the cultures 10 times and incubated them for an additional 20 minutes at 37 °C. We transferred the cultures to a 1.5 mL microfuge tube and spun them at 500 × g for 5 minutes at room temperature (RT) in a Sorvall tabletop centrifuge. We resuspended the cell pellets in 0.5 mL of 0.25% trypsin-EDTA (Gibco 25200-056) and incubated them for 15 minutes at 37 °C. We then triturated the cultures and incubated them for an additional 15 minutes. We added 0.5 mL of media, triturated, and spun the samples at 500 × g for 5 minutes at RT. We washed the cell pellets once with 1 mL PBS and subsequently used them in the protocols described below. While undifferentiated cells detach in the presence of trypsin-EDTA alone, differentiated cells require incubation with collagenase as outlined; therefore, we treated cells in all states the same for continuity.

### hFOB ATAC-seq and RNA-seq

We performed ATAC-seq on osteoblast-differentiated hFOB cells at 2, 4, and 10 days post-induction, as well as on undifferentiated cells, using an ATAC-seq kit (Active Motif 53150) and following the manufacturer’s recommended protocol. Briefly, on the indicated days, we detached cells from the culture plates as outlined previously, isolated nuclei from 100,000 cells per sample, and subsequently tagmented, purified, and stored the nuclear DNA at −20 °C until all samples had been processed. We repeated this sample preparation scheme three additional times, resulting in a total of 16 samples overall from four separate experiments using four different hFOB passages. We performed PCR amplification of tagmented DNA and constructed Illumina sequencing libraries for all samples en masse. We pooled the libraries and paired-end sequenced them to a length of 150 bp per end on a single lane of an Illumina HiSeqX at a depth of approximately 50 million paired-end reads per sample. We uniformly processed ATAC-seq reads, aligned them to hg38, and called peaks jointly across the four biological replicates at each time point using Genrich (github.com/jsh58/Genrich) in ATAC-seq mode with PCR duplicates removed. Replicates were combined to identify significantly accessible regions, which we reported in ENCODE narrowPeak format. We assessed data quality and reproducibility across biological replicates and time points using the nf-core/atacseq pipeline (v2.12.2), which performed read trimming, alignment to hg38, duplicate marking and filtering, and narrow peak calling with MACS2 ^64^. We performed principal component analysis (PCA) across all 16 samples to assess clustering of replicates within and between time points (**Extended Data Fig. 2a**).

Short-read bulk RNA-seq data were previously generated from hFOB cells at six time points of osteogenic differentiation (days 0, 2, 4, 6, 8, and 10), with three biological replicates per time point, as described in Abood *et al*. ^30^. We aligned reads to the human reference genome (hg38), quantified gene expression, and performed differential expression analyses across the time course using DESeq2 ^65^. Long-read RNA-seq data were previously generated from hFOB cells at four time points (days 0, 2, 4, and 10), with three biological replicates per time point, as described in Abood *et al*. ^31^. We assessed differential gene and transcript expression, and differential isoform usage, across the differentiation time course using tappAS ^66^.

### hFOB CRISPR inhibition and activation cell lines

We generated hFOB cell lines that constitutively express either the CRISPRi vector KRAB-dCas9-MeCP2 (Addgene 170068) ^6^ or the CRISPRa vector dCas9-VPR ^34,35^ by packaging the respective plasmids into lentiviral particles via polyethylenimine-mediated transfection of HEK293 cells along with the packaging plasmids pMD2.G and psPAX2 (Addgene 12259, 12260), following the manufacturer’s recommendations (Polysciences 23966). We collected the virus and concentrated it 20-fold from media 3 days after transfection (following the manufacturer’s protocol; Alstem VC100) and used it to transduce hFOB cells with varying amounts of virus in growth media (minus antibiotics) containing 8 µg/mL polybrene, following the manufacturer’s recommendations (Millipore TR-1003). We passaged transduced cells in growth media containing 5 µg/mL blasticidin for at least 2 weeks prior to DNA and RNA isolation and analyzed them via qPCR for dCas9 copy number and expression level (see **Supplementary Table 3b** for description of primers used). We selected the cell population with the highest relative copy number and expression that did not adversely affect growth or osteoblast differentiation potential for use in the experimental studies described herein.

### Genome-wide association study of estimated BMD

We used UK Biobank data upon ethical approval from the Northwest Multi-Centre Research Ethics Committee, and the UK Biobank informed consent from all participants before participation. We analyzed genome-wide association study (GWAS) summary statistics of BMD estimated from quantitative heel ultrasounds in 426,824 individuals that we generated previously ^2^. We performed this GWAS using a linear mixed non-infinitesimal model, adjusting for age, sex, genotyping array, assessment centre, and ancestry-informative principal components 1–20 to account for population structure and cryptic relatedness. This GWAS identified 515 significant loci across chromosomes 1 through 22 with 1,097 conditionally independent signals, which we detected using GCTA-COJO ^67^ (*p*_COJO_ < 6.6 × 10⁻⁹).

### Statistical fine-mapping GWAS variants

We performed statistical fine-mapping of BMD GWAS loci to identify 95% credible sets containing likely causal variants using the Sum of Single Effects (SuSiE) model ^36^. SuSiE uses GWAS summary statistics and population-matched linkage disequilibrium (LD) correlation matrices to identify independent credible sets within a GWAS locus. We generated LD correlation matrices for all 515 significant loci using a subset of 50,000 individuals from the UK Biobank, defining each locus as spanning ± 500 kb around all conditionally independent signals within that locus. We then fit SuSiE models for each locus, testing for up to 25 credible sets per locus. We successfully fine-mapped 499 loci, with 16 failing to identify any credible sets, resulting in 1,353 distinct credible sets and 27,099 fine-mapped variants. We removed variants with less than 1% posterior inclusion probability, and our final set of fine-mapped variants included 17,003 variants from 1,350 credible sets across 499 loci.

### BMD STING-seq gRNA library design

We annotated protein coding fine-mapped BMD variants using the Variant Effect Predictor ^68^. We then integrated hFOB epigenomic data from days 0, 2, and 4 of differentiation. To identify candidate *cis*-regulatory elements (cCREs) for CRISPRi targeting, we intersected noncoding fine-mapped BMD variants with hFOB ATAC-seq peaks or selected variants located at least 1 kb from an hFOB ATAC-seq peak summit. We prioritized cCREs where one of the three closest genes was a transcription factor ^37^ expressed at a level of at least 0.5 TPM in hFOB long-read RNA-seq data ^30^. This process resulted in a set of 140 BMD GWAS loci with 202 distinct credible sets and 570 fine-mapped noncoding variants. We then identified a representative variant for those where multiple variants were within 1 kb of each other, as CRISPRi has an approximate 1 kb effect radius ^6^, resulting in 393 variants corresponding to 140 loci. We designed gRNAs to target within 150 bp of a variant and used FlashFry v1.10.0 ^69^ to retain gRNAs with the lowest predicted off-target activity, as estimated by the Hsu-Scott score ^39^. We obtained a final list of 361 variants, targeted by 3 gRNAs each, corresponding to 138 loci.

We designed gRNAs targeting the transcription start sites (TSS) of TFs, or genes annotated as possibly functioning as TFs ^6,37^, bearing fine-mapped protein coding BMD GWAS variants, that were expressed at a level of at least 0.5 TPM in hFOB long-read RNA-seq data ^30^. We identified variants with predicted deleterious effects on protein function if they were annotated as MODIFIER or HIGH impact variants. Given that genes may express multiple isoforms, we designed gRNAs to target each expressed isoform’s TSS if they were over 1 kb apart. Upon applying the same off-targeting criteria above, we obtained a final list of 59 TSS’s, targeted by 3 gRNAs each, corresponding to 23 genes (5 not captured in hFOB scRNA-seq, therefore we report 18 in our results). We also included 30 non-targeting gRNAs from the GeCKOv2 library ^43^ as negative controls and 22 *CD55* TSS-targeting gRNAs as multiplicity-of-infection (MOI) controls, as we have described previously ^6^ and will further describe below.

### High MOI single-cell CRISPR screens

We synthesized the BMD STING-seq gRNA library oligos and cloned them into the plasmid lentiGuideFE-Puro (Addgene 170069) immediately 3′ of the U6 promoter using the Gibson assembly reaction, following the manufacturer’s recommendations (NEB E2621S). We amplified the plasmid library by transfecting electrocompetent Stbl4 E. coli cells and then isolated plasmid DNA (ThermoFisher 11635018, K21007). We verified individual gRNA representation by amplicon sequencing prior to packaging the plasmid library into lentiviral particles via polyethylenimine-mediated transfection of HEK293 cells along with the packaging plasmids pMD2.G and psPAX2 (Addgene 12259, 12260), following the manufacturer’s recommendations. We concentrated the virus 20-fold from media 3 days after transfection (Alstem VC100) and used it to transduce CRISPRi hFOB cells for our BMD STING-seq experiments. We transduced CRISPRi hFOB cells with the lentiviral library, selected them in growth media containing 3 µg/mL puromycin (Gibco A11138-03) starting 2 days after transduction, and continued selection for 5 days. At that point, we either plated cells for the time points collected in our full single-cell experiment (described below) or collected them for the pilot single-cell experiment. For the full experiment, we plated cells contributing to the day 4 sample 7 days after transduction, subjected them to osteogenic growth conditions 2 days after plating, and harvested them 4 days later. For the day 2 sample, we plated cells 9 days after transduction and treated them as outlined for the day 4 sample, except we harvested them 2 days after induction of osteogenic differentiation conditions. For the undifferentiated (day 0) sample, we plated cells 11 days after transduction and harvested them 2 days later. The transduced cells had an average MOI of 10. We predetermined the amount of virus library needed to achieve this MOI by qPCR analysis of the relative copy number of the puromycin gene, using DNA collected from cells transduced with varying amounts of the BMD STING-seq lentiviral library.

We performed single-cell RNA-seq using 10X Genomics Chromium Next GEM Single Cell 5′ Reagent Kits v2 (dual index) on cells detached from culture plates. We labeled cells from each state with different hashtag antibodies targeting cell surface proteins (BioLegend TotalSeq 394661, 394663, 394665, 394667) following the manufacturer’s recommendations. We then pooled equal numbers of cells from each state into a single sample and aliquoted them into as many reactions as needed to capture approximately 200,000 cells in total. We used 2 microfluidics channels for the pilot experiment and 6 channels for the full experiment. We constructed sequencing libraries for the three different barcoded nucleic acids for each reaction according to 10X Genomics protocols and pooled them disproportionately so that mRNA-derived cDNA libraries represented ∼90% of the sample, with gRNA and hashtag libraries comprising the remaining 10%. We sequenced the pooled library to a depth of approximately 20,000 mRNA/cDNA reads per cell on an Illumina NextSeq 500 sequencing platform for the pilot experiment and on an Illumina NovaSeq X for the full experiment.

### Multi-modal single-cell data processing

We generated UMI count matrices for all single-cell sequencing libraries using 10X Cell Ranger v8.0.1. We produced cDNA UMI count matrices aligned to hg38 (refdata-gex-GRCh38-2024-A), gRNA UMI count matrices, and hashtag UMI count matrices for the pilot and full experiments. We analyzed the UMI count matrices in R v4.5.2 using Seurat v5.3.1. We manually inspected the distributions of cDNA, gRNA, and hashtag UMIs for each microfluidics channel and set custom thresholds to remove outliers for total cDNA count, unique genes detected, mitochondrial percentage, total gRNA count, unique gRNAs detected, total hashtag count, and unique HTOs detected. We grouped cells generated on different microfluidics channels for the pilot and full experiments only after quality control checks have been performed on channels individually.

### Differential gene expression testing

We utilized the processed UMI count matrices for gene expression and gRNA expression, along with accompanying single-cell metadata as covariates in model fitting. For each cCRE, we defined a list of genes within ±1 Mb to test for differential expression in *cis*. We then tested for differential outcomes within the SCEPTRE framework ^47^, adjusting for the following single-cell covariates in expression tests: total gene expression UMIs, unique genes, total gRNA expression UMIs, unique gRNAs, percentage of mitochondrial genes, microfluidics channel, UMI count of dCas9 expression, and the first 25 gene expression principal components. We used SCEPTRE for differential gene expression testing, as we previously demonstrated SCEPTRE is optimal for high MOI single-cell CRISPR screens. To test for differential expression in trans, we defined for each set of gRNAs a list of all genes detected in at least 5% of cells and repeated the test above. We tested non-targeting gRNAs against all genes used in the *cis* and *trans* settings and randomly sampled them to match the number of *cis* and *trans* tests displayed on QQ-plots. We use the same covariates for targeting gRNAs and we do for non-targeting gRNAs. We do not perform differential expression testing of *CD55*-targeting gRNAs as these were solely used to titrate for a high-MOI experiment. To determine significance for multiple hypotheses (genes) tested in *cis*, we adjusted SCEPTRE *p*-values using the Benjamini-Hochberg procedure. For STING-seq analyses, we reported *cis*-target genes as significant at a 5% FDR (Benjamini–Hochberg adjusted *p*_SCEPTRE_ < 0.05) and defined *trans*-target genes as those significant at a stricter 1% FDR.

### Consensus non-negative matrix factorization (cNMF)

To capture gene programs within our BMD STING-seq data, we filtered for genes expressed in at least 10% of cells, resulting in a set of 9,604 genes, from all time points. We performed cNMF (100 iterations) to decompose the expression matrix into two components: a usage matrix of cellular program activity and a spectra matrix of gene weights. One component is a gene-weight matrix representing co-regulated gene programs, while the other constitutes a program-usage matrix reflecting the relative contribution of each program to the transcriptome of individual cells. We utilized a default density threshold of 0.5 to remove outlier programs and tested *K* from 35 to 80 (step size 5; step size 1 for *K* = 40–50). We selected the optimal number of programs (*K*=45) based on the highest stability, measured by the Euclidean distance silhouette score, relative to the consensus solution’s Frobenius error. Each program was defined using the spectra matrix, specifically as the top 300 genes ranked by their cNMF-derived *Z*-score regression coefficients (**Supplementary Table 5c**).

### Identifying programs associated with targeted variants

To assess the association between perturbations and the 45 gene programs, we constructed a binary perturbation matrix. We collapsed individual gRNAs targeting the same gene or SNP into single features; to ensure comparable noise properties, non-targeting controls were randomly aggregated into groups of three. We then modeled the activity of each program (usage matrix) as a function of perturbation status using linear regression. The model was adjusted for technical covariates (including detected genes/UMIs for both cDNA and gRNA capture, and mitochondrial percentage), experimental batch, and top 25 principal components; (**Supplementary Table 2e**). Significant associations were identified at a 1% FDR using the Benjamini–Hochberg adjustment.

### hFOB whole-genome sequencing and allele-specific open chromatin

Genomic DNA was extracted from hFOB cells using standard methods. A total of 500 ng of high-quality genomic DNA was submitted to Psomagen (Rockville, MD, USA) for whole-genome sequencing. Sequencing libraries were prepared using the TruSeq Nano DNA Library Prep Kit (Illumina) according to the manufacturer’s protocol. Libraries were sequenced on an Illumina NovaSeq 6000 S4 platform using 150 bp paired-end reads, generating ∼185 Gb of raw sequence data, corresponding to a mean genome coverage of ∼50x. Primary bioinformatics processing was performed by Psomagen as part of their standard WGS analysis pipeline. This included sequencing quality control, read alignment to the human reference genome, and germline variant calling. Processed alignment files (BAM) and variant call files (VCF), along with raw sequencing data (FASTQ), were returned via secure transfer for downstream analyses.

We determined variants mapping to the targeted screens were heterozygous in hFOB cells by querying their genomic positions in the hFOB whole-genome. We then examined the genotype annotations in the VCF to classify each variant as heterozygous, homozygous reference, or homozygous alternate.

We corrected for mapping bias using WASP ^70^ then obtained allele-specific read counts using GATK ASEReadCounter ^71^. In R, we merged counts across replicates to increase read depth, filtered out variants with fewer than 10 reads, performed a binomial test, and applied adjusted p-values for multiple testing using the Benjamini-Hochberg method.

### Arrayed CRISPRi and CRISPRa validation experiments

We designed CRISPR gRNAs targeting open chromatin regions (promoters and enhancer regions) as described previously or using the UCSC Genome Browser “CRISPR targets” track. We synthesized oligonucleotides containing the chosen sequences with the addition at the 5′ end of the reverse complement of the 4-nucleotide overhang generated after digestion of the lentiGuideFE-Puro plasmid (Addgene 170069) with the BsmB1 restriction enzyme (NEB R0739L; **Supplementary Table 3a**). We validated plasmids for identity and accuracy by sequencing and subsequently packaged them into viruses as outlined above. We transduced the viruses into hFOB CRISPRi and/or hFOB CRISPRa cells as described previously. The results reported in this study are based on four such experiments.

### Replication of CRISPRi effects in hMSCs

Primary human mesenchymal stem cells (hMSCs) were derived from bone recovered from the acetabular reamings of a male patient collected during total hip arthroplasty. Specimen collection was approved by the University of Virginia Institutional Review Board for Health Sciences Research (IRB-HSR HSR200330) and written informed consent was obtained from the donor. Briefly, washed reamings from an individual patient were sparsely distributed in a 10 cm cell culture plate that contained 20 mL of growth media. Media was replaced every three days and reamings were removed after 10–14 days. Media changes continued as outlined until the plate was 80–90% confluent at which time cells were detached with 0.25% trypsin/EDTA washed and frozen in one or more aliquots (5–7 days after removal of the reamings). For shRNA knockdown experiments, a vial of the MSCs were thawed, washed, dispersed in a cell 15 cm culture plate and allowed to grow to 80–90% confluence, changing media every three days, if necessary, at which time cells were detached, washed, counted and plated for transduction the following day.

Lentiviral production and transduction followed the procedures described for the arrayed hFOB validation experiments, except that hMSCs were co-transduced with lentiviruses encoding the individual gRNA and KRAB-dCas9-MeCP2. RNA was collected 3–10 days after transduction. RNA isolation, cDNA synthesis, and RT-qPCR followed the procedures described in the corresponding subsections below, with transcript abundance normalized to CHMP2A and expressed relative to matched non-targeting controls using the 2^−ΔΔCt method. Each guide– transcript comparison was evaluated in three or four independent experiments using cells from the same donor.

### Validation of *trans*-regulatory networks in hMSCs

To validate trans-regulatory networks identified by STING-seq, we performed targeted gene knockdown followed by bulk RNA-seq in primary human MSCs from the same donor described above. Lentiviral shRNAs targeting *TRIP11* (TRCN0000022019 and TRCN0000022021; Sigma-Mission) and *DAP3* (TRCN0000229949 and TRCN0000229951; Sigma-Mission), together with an empty vector control (SHC001; Sigma-Mission), were packaged into lentiviral particles and used to transduce hMSCs as described above. In parallel, *TRIP11* was targeted using lentiviral CRISPRi. RNA was collected at days 6 and 10 post-transduction using the RNeasy Mini Kit (Qiagen) and submitted to Psomagen (Rockville, MD) for bulk RNA-seq. Raw reads were processed using nf-core/rnaseq (v3.14.0) with Nextflow (v24.04.4). Transcript abundance was quantified using Salmon against GRCh38 (GENCODE v43) and aggregated to gene-level counts using tximport in R (v4.5.0). Genes with fewer than 10 total counts were excluded. Differential expression was assessed separately at each time point using DESeq2 (v1.52.0), adjusted by biological replicate. *P*-values from Wald tests were adjusted using the Benjamini-Hochberg method. Predefined *TRIP11* and *DAP3*–*YY1AP1* network genes were also analyzed separately, with FDR calculated within each network. To assess network conservation, we compared gene-level log_2_ fold-changes between the hMSC bulk RNA-seq and hFOB STING-seq experiments. Because effect sizes are not directly comparable between the two assays, concordance was assessed based on the direction of effect. At each time point, the proportion of genes with concordant effects was tested against the 50% expected by chance using a one-sided exact binomial test, both across all overlapping network genes and among genes significant in the hMSC bulk RNA-seq analysis.

### Examining CRE–gene pairs for colocalized BMD GWAS variants and GTEx eQTLs

We performed colocalization analyses between BMD GWAS signals ^26^ and expression quantitative trait loci (eQTLs) using fastENLOC v3.1 ^72^ applied to GTEx v8 data ^73^ (**Supplementary Table 4**). These GWAS and eQTL fine mapping results were integrated using fastENLOC to compute regional colocalization probabilities (RCPs) between BMD loci and gene expression, and genes with RCP > 0.1 were considered to show evidence of colocalization. For each colocalized gene, we annotated the GWAS posterior inclusion probability (PIP) of the colocalization driving variant and summarized tissue specificity by reporting the number of GTEx tissues in which colocalization was observed under two criteria: a filtered set requiring that at least one colocalization driving GWAS variant reached genome wide significance for BMD, and an unfiltered set including all eQTLs with RCP > 0.1 irrespective of GWAS significance. We additionally reported cases in which a given variant drove colocalization with multiple eQTLs (distinct target genes), together with tissue counts under both definitions, and when a gene showed evidence of colocalization but the queried variant itself was not the primary driver, we reported all variants inferred to drive the signal, sorted by GWAS PIP and annotated by rs ID, hg38 genomic coordinate, reference and alternate alleles, GWAS PIP, and the number of supporting tissues.

### Targeting *DAP3* and *YY1AP1* with siRNAs

We transfected hFOB cells used in siRNA knockdown experiments within 5 days of thawing from liquid nitrogen storage. In a typical experiment, we seeded 13 wells for each siRNA treatment (three siRNAs per gene plus a non-targeting control; **Supplementary Table 6i**) as well as a null transfection control, for a total of 104 wells. We used four wells of each treatment to determine the amount of mineral formed at various differentiation days (7–10), and the remaining wells for RNA isolation at various time points during the experiment. We seeded cells at 75,000 cells per well in a 24-well plate and transfected them within 24 hours using siRNAs and lipofectamine LTX reagent (Invitrogen 15338100) following the manufacturer’s recommended procedure with minor modifications. Briefly, we mixed 2.5 µL lipofectamine LTX with 37.5 µL pre-warmed Opti-MEM (Gibco 31985-070) per well of a 24-well plate. In parallel, we mixed 1 µL of 5 µM siRNA with 37.5 µL pre-warmed Opti-MEM per well. After incubating each mixture for at least 5 minutes at RT, we combined them, mixed, and incubated for a minimum of 15 minutes at RT. We then diluted the ∼75 µL lipofectamine and siRNA mixture into 500 µL pre-warmed Opti-MEM per well, mixed, and applied it to cells after removing the growth media. We placed the cells back in the incubator (34 °C) for 5–7 hours, after which we removed the Opti-MEM transfection mix, washed the cells with pre-warmed Dulbecco’s Phosphate Buffered Saline (DPBS, Gibco 14190-144), and replaced it with 500 µL per well of pre-warmed growth media. The results reported in this study are based on three such experiments (**Supplementary Table 6k–m**).

### RNA isolation and cDNA preparation

We isolated RNA using the RNeasy Mini Kit (Qiagen 74106) following the manufacturer’s protocol. Briefly, we removed media from cells, washed them three times with 1 mL per well of DPBS (Gibco 14190-144), and lysed them in 350 µL per well of RLT buffer containing 40 mM dithiothreitol (DTT) with mild shaking for 10 minutes. We transferred the lysates to microcentrifuge tubes and stored them at −80 °C until processing. At the time of processing, we isolated RNA from thawed lysates exactly as outlined in the manufacturer’s protocol. We immediately DNase-treated the RNA using Applied Biosystems TURBO DNA-free Kit (Invitrogen AM1907) following the manufacturer’s protocol and stored DNase-treated samples at −80 °C. We prepared cDNA from thawed samples by first determining RNA concentration using a Qubit 4 fluorometer and the RNA HS or BR assay kits (ThermoFisher Q33226, Q32855, Q10211) according to the manufacturer’s protocol. We synthesized random-primed cDNA from 1–2 µg of DNase-treated RNA using a High Capacity Reverse Transcription Kit (Applied Biosystems 4368813) following the manufacturer’s protocol.

### Measuring CRISPR inhibition, activation or knockdown with qPCR

We determined the relative abundance of different transcripts in duplicate for each sample or treatment from four separate experiments at least 7 days after transduction and 2–10 days after siRNA transfection. Each qPCR reaction contained 10 ng cDNA, 800 nM of each primer (**Supplementary Table 3a**), and 0.5× PowerUp SYBR Green Master Mix (Applied Biosystems 100029285) in a 10 µL reaction volume. We amplified reactions on a QuantStudio 5 Real-Time PCR System (Applied Biosystems A28135) under the following cycling conditions: 50 °C for 2 minutes, 95 °C for 2 minutes; followed by 40 cycles of 95 °C for 1 second and 60 °C for 30 seconds; and a melt curve of 95 °C for 1 second, 60 °C for 20 seconds, and 95 °C for 1 second. We determined relative quantification of qPCR products using the 2^−ΔΔC(t)^ method ^74^, using the C(t) of *CHMP2A* as a reference, as we previously confirmed that *CHMP2A* expression does not change during osteoblast differentiation in the hFOB cell line ^31^. Briefly, we calculated the cycle number at which half of the final product amount was produced (Cq/C(t)) following the manufacturer’s recommendations. For each sample and primer pair, we subtracted the C(t) value of *CHMP2A* for the same sample to obtain ΔC(t). We then subtracted the ΔC(t) of the non-targeting control sample from the ΔC(t) of the gRNA transduced or siRNA transfected sample for that primer pair to obtain ΔΔC(t). We reported values as 2 raised to the power of negative ΔΔC(t).

### Crystal violet staining for cell viability

We performed cell number assays using hFOB CRISPRi cells transduced in an arrayed fashion with two gRNAs targeting the DAP3–YY1AP1 pCRE and a non-targeting gRNA. One gRNA was from the BMD STING-seq library (pCRE-1), and the other was designed with the UCSC Genome Browser “CRISPR targets” track (pCRE-2). We seeded each cell population into 24-well culture plates at a density of 10,000 cells per well, in four cells each, and grew them in MEM alpha supplemented with 10% FBS for 3 days before treating them with osteogenic medium for 7 or 10 days. At the end of the treatment period, we removed the culture medium and washed wells once with PBS to remove residual medium and dead cells. We fixed cells with 4% paraformaldehyde (PFA) for 15 minutes at room temperature, washed each well twice with distilled water, and air-dried the plates completely. We stained fixed cells with 0.1% crystal violet (Sigma-Aldrich C6158) prepared in PBS for 20 minutes at room temperature with gentle rocking. After staining, we removed excess dye by thoroughly rinsing the plates with tap water until the wash ran clear and then air-dried the plates completely at room temperature. We captured images using a CanoScan 9000F (Canon). For quantification, we solubilized bound crystal violet by adding 400 µL of 1% acetic acid (v/v) to each well and incubating for 10–15 minutes with gentle shaking to ensure complete dye dissolution. We measured absorbance at 590 nm using a SmartSpec Plus Spectrophotometer (BioRad). The optical density (OD) values correlated with the number of adherent cells, serving as an indirect measure of cell proliferation or viability. Quantification was performed from three independent experiments.

### Sirius Red staining for measuring total collagen content

We seeded hFOB CRISPRi cells transduced with the NT, pCRE-1, and pCRE-2 gRNAs introduced above into 24-well culture plates at a density of 10,000 cells per well, 4 wells each, and grew them in MEM alpha supplemented with 10% FBS for 3 days before treating them with osteogenic medium for 7 or 10 days. Following treatment, we aspirated the medium and gently washed cells twice with PBS to remove residual serum proteins. We fixed cells with Kahle’s solution, containing 26% ethanol, 3.7% formaldehyde, and 2% glacial acetic acid, at room temperature for 15 minutes. After fixation, we washed wells thoroughly with running tap water until the yellow color was completely removed. We stained fixed cells with 0.1% (w/v) Direct Red 80 (Sigma-Aldrich 365548) dissolved in 1% acetic acid for 1 hour at room temperature with gentle shaking. After staining, we washed wells with 400 µL of 0.1 M HCl and then eluted collagen-bound dye using 400 µL per well of 0.1 M NaOH. Before dye elution, we air-dried plates completely at room temperature and captured images using a CanoScan 9000F (Canon). We measured the absorbance of the eluted dye at 540 nm using a SmartSpec Plus Spectrophotometer (BioRad). OD values correlated with collagen content in each sample. We normalized collagen content to cell number using crystal violet staining. Quantification was performed from three independent experiments.

### Seahorse XFe96 assay for measuring mitochondrial respiration and bioenergetics

We seeded hFOB CRISPRi cells transduced with the NT, pCRE-1, and pCRE-2 gRNAs into Seahorse XFe96 cell culture microplates (Agilent) at a density of 8,000 cells per well in 80 µL of complete growth medium and incubated plates overnight at 37 °C and 5% CO₂ to allow cell attachment. The following day, we changed the medium to osteogenic medium (OM) and incubated cells for 48 hours. Twenty-four hours prior to the measurement, we hydrated the XFe96 sensor cartridge by adding 200 µL per well of Seahorse XF Calibrant (Agilent) to the accompanying utility plate and incubated it overnight at 37 °C in a non-CO₂ incubator to equilibrate the sensors. On the day of the assay, we replaced the culture medium with Seahorse XF DMEM (Agilent) supplemented with 10 mM glucose, 1 mM sodium pyruvate, and 2 mM L-glutamine. Each well was washed twice with 180 µL of assay medium and then incubated with 180 µL of fresh assay medium at 37 °C in a non-CO₂ incubator for 60 minutes to allow temperature and pH equilibration. We prepared 10x working concentrations of mitochondrial stress compounds— oligomycin (1 µM final, ATP synthase inhibitor), FCCP (1 µM final, mitochondrial uncoupler), and rotenone plus antimycin A (0.5 µM final each, Complex I and III inhibitors)—in assay medium and loaded them into injection ports A–C of the hydrated sensor cartridge. We calibrated the hydrated cartridge using the Seahorse Wave software (Agilent), after which we loaded the cell plate and performed the Mitochondrial Stress Test according to the manufacturer’s protocol. We measured baseline oxygen consumption rate (OCR) and extracellular acidification rate (ECAR) prior to compound injections. Sequential injections of oligomycin, FCCP, and rotenone/antimycin A were used to dissect key components of mitochondrial respiration. OCR values were normalized to cell number using post-assay nuclear staining with DAPI.

### Confocal microscopy of mitochondrial morphology

We seeded hFOB CRISPRi cells transduced with the NT, pCRE-1, and pCRE-2 gRNAs onto sterile glass coverslips at a density of 50,000 cells per well in 6-well plates and cultured them overnight in complete growth medium at 37 °C and 5% CO₂ to allow cell attachment. The following day, we changed the medium to OM and incubated cells for 48 hours. After 48 hours, we treated cells with 100 nM MitoTracker Red CMXRos (ThermoFisher M7512) for 30 minutes at 37 °C under standard culture conditions, followed by two gentle washes with pre-warmed PBS to remove excess dye. We fixed cells with 4% PFA for 15 minutes, washed them twice with PBS, and mounted coverslips using ProLong Gold Antifade Mountant with DAPI (Invitrogen P36971). We acquired images on a Zeiss LSM 880 Axio Inverted laser-scanning confocal microscope (Plan-Apochromat 63×/1.40 oil). Raw confocal images were processed in Fiji/ImageJ (NIH), and mitochondrial morphology and network parameters were analyzed using the MiNA (Mitochondrial Network Analysis) plugin in ImageJ. Quantification was performed on at least 50–60 cells per condition from three independent experiments.

## Extended Data Figures

**Extended Data Fig. 1.**
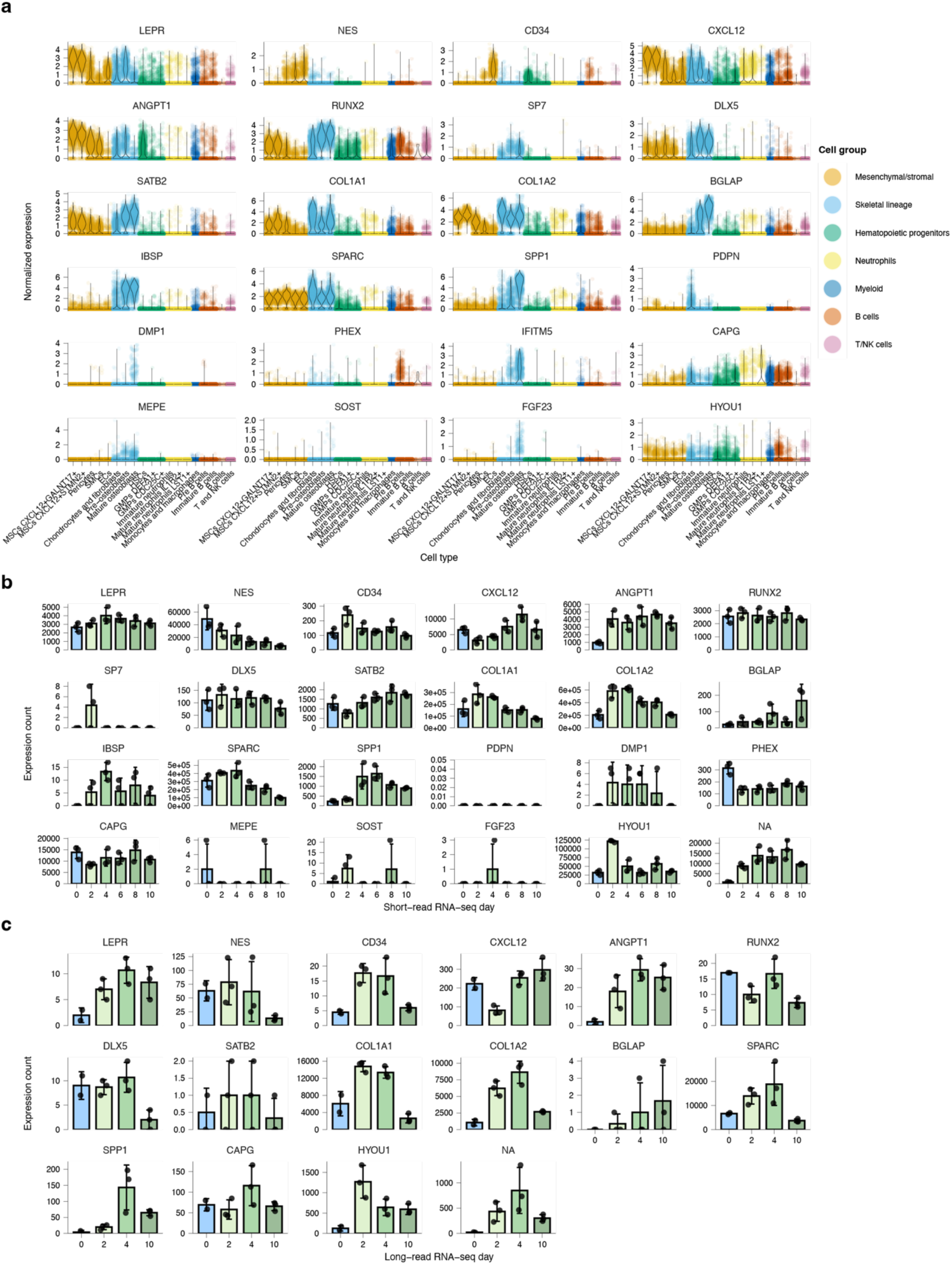
hFOB cells recapitulate the osteoblast marker gene expression patterns of a human bone single-cell atlas across both short- and long-read RNA-seq time courses. **a,** Violin plots showing normalized expression of osteoblast, osteocyte, and stromal/hematopoietic marker genes across cell types from a published single-cell RNA-seq atlas of the human bone marrow niche (Chai *et al*. ^26^), color-coded by cell group (mesenchymal/stromal, skeletal lineage, hematopoietic progenitors, neutrophils, myeloid, B cells, and T/NK cells). **b,** Expression of the same marker genes in hFOB cells across the short-read RNA-seq osteogenic differentiation time course (days 0, 2, 4, 6, 8, and 10). Bars show mean expression count ± standard deviation, with points representing individual replicates. **c,** Expression of the same marker genes in hFOB cells across the long-read RNA-seq osteogenic differentiation time course (days 0, 2, 4, and 10). Not all marker genes shown in **a** were detected at sufficient levels for quantification in the short- or long-read datasets.

**Extended Data Fig. 2.**
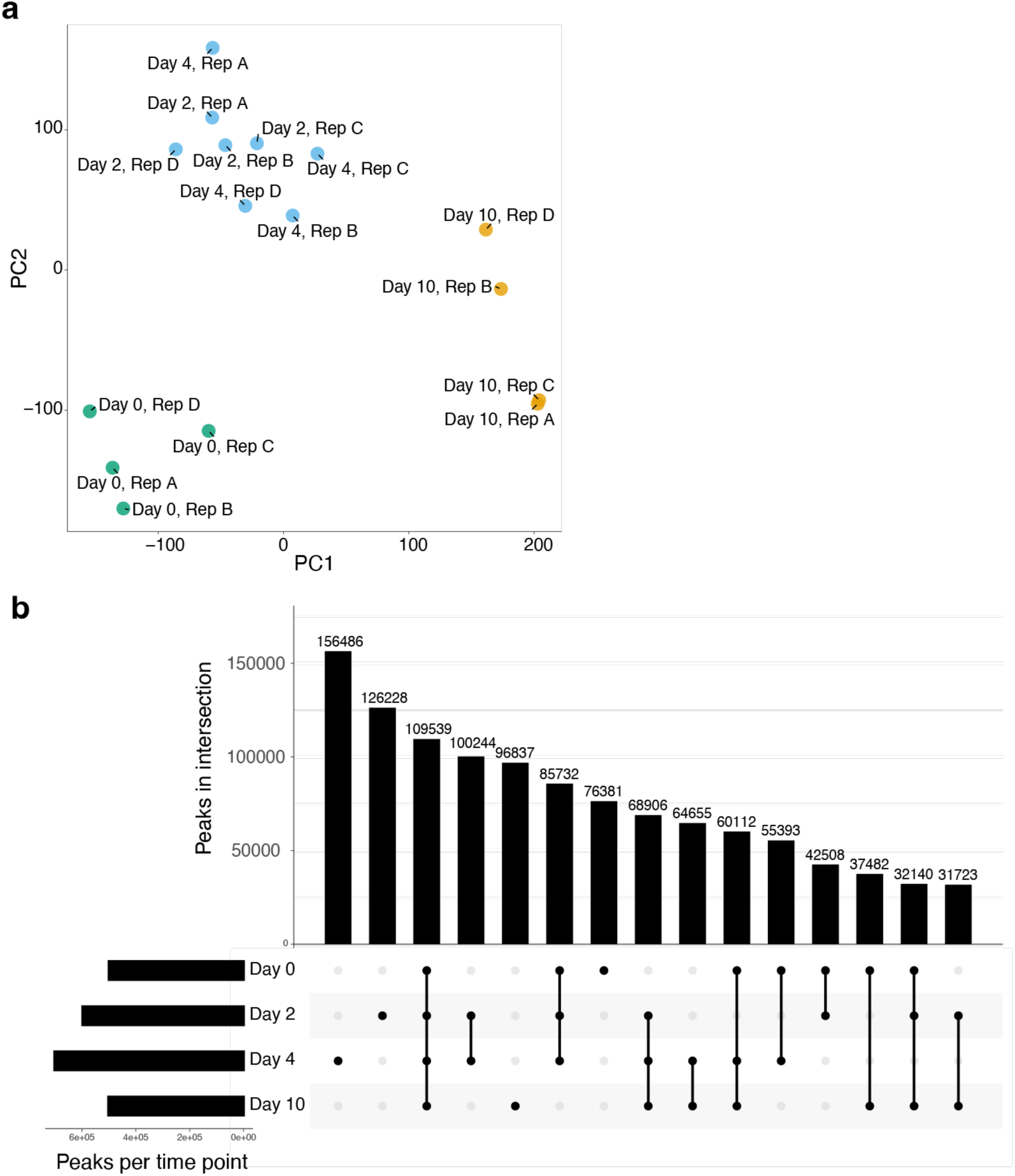
ATAC-seq reproducibility and chromatin accessibility dynamics during hFOB osteogenic differentiation. **a,** Principal component analysis of ATAC-seq peak accessibility across four biological replicates (Rep A–D) at days 0, 2, 4, and 10 of osteogenic differentiation. Replicates clustered by differentiation stage. Day 0 and day 10 samples formed distinct clusters, whereas days 2 and 4 clustered together, consistent with similar early osteoblast-like states. **b,** UpSet plot showing overlap of ATAC-seq peaks across differentiation time points. Horizontal bars indicate the total number of peaks identified at each time point, and vertical bars indicate the number of peaks unique to or shared among the indicated time points. A core set of 109,539 peaks was shared across all four time points. Day 4 contained the largest number of time point-specific peaks (156,486), whereas day 10 contained the fewest accessible regions overall, consistent with reduced chromatin accessibility in more mature, mineralizing osteoblast-like cells.

**Extended Data Fig. 3.**
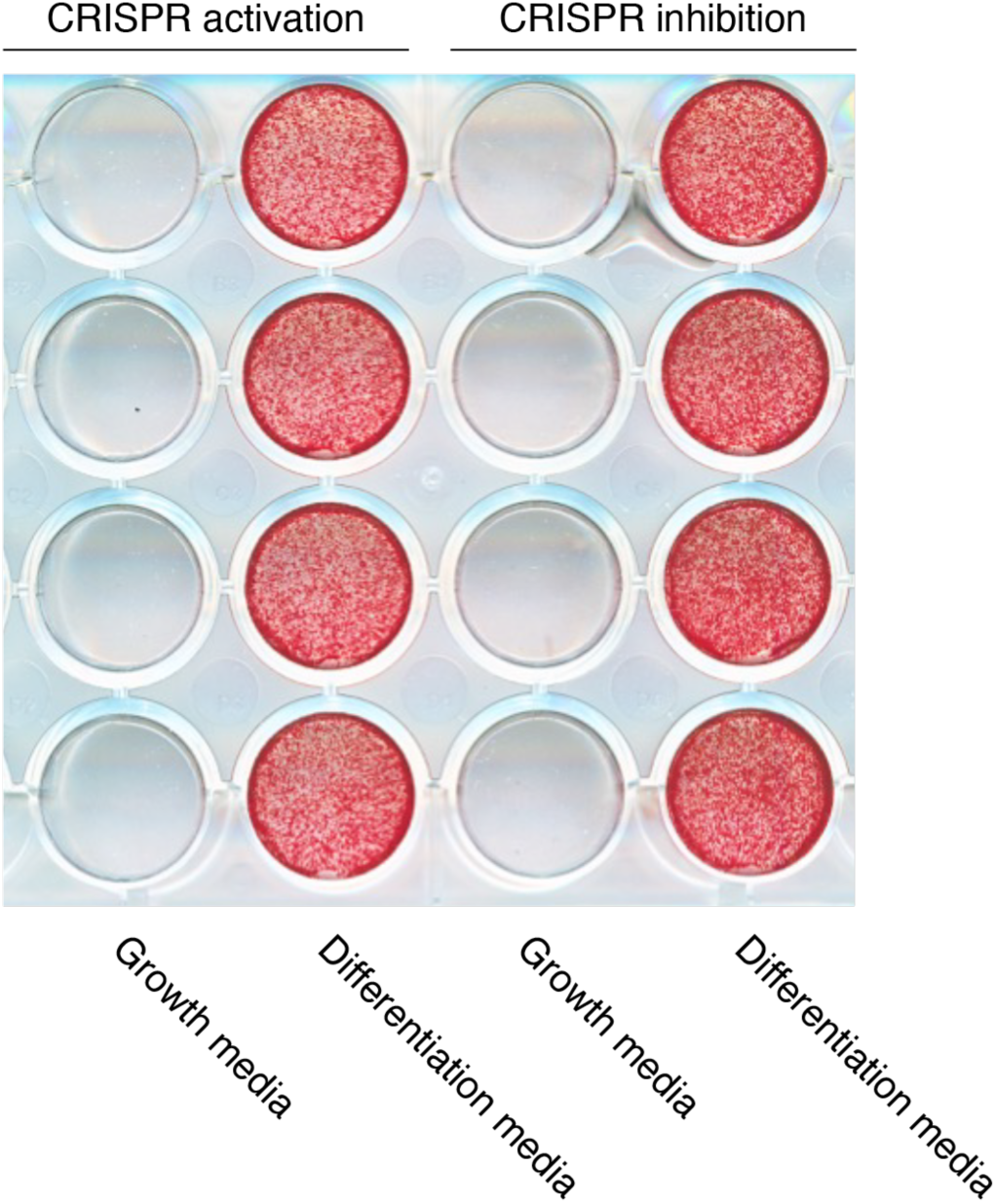
hFOB cells expressing the CRISPRi and CRISPRa machinery retain the capacity to mineralize. Alizarin Red staining of hFOB cells constitutively expressing CRISPRi (KRAB-dCas9-MeCP2) or CRISPRa (dCas9-VPR) at day 10 of culture in growth media compared to day 10 of culture post-osteogenic induction in differentiation media. Four replicate wells are shown per CRISPR effector.

**Extended Data Fig. 4.**
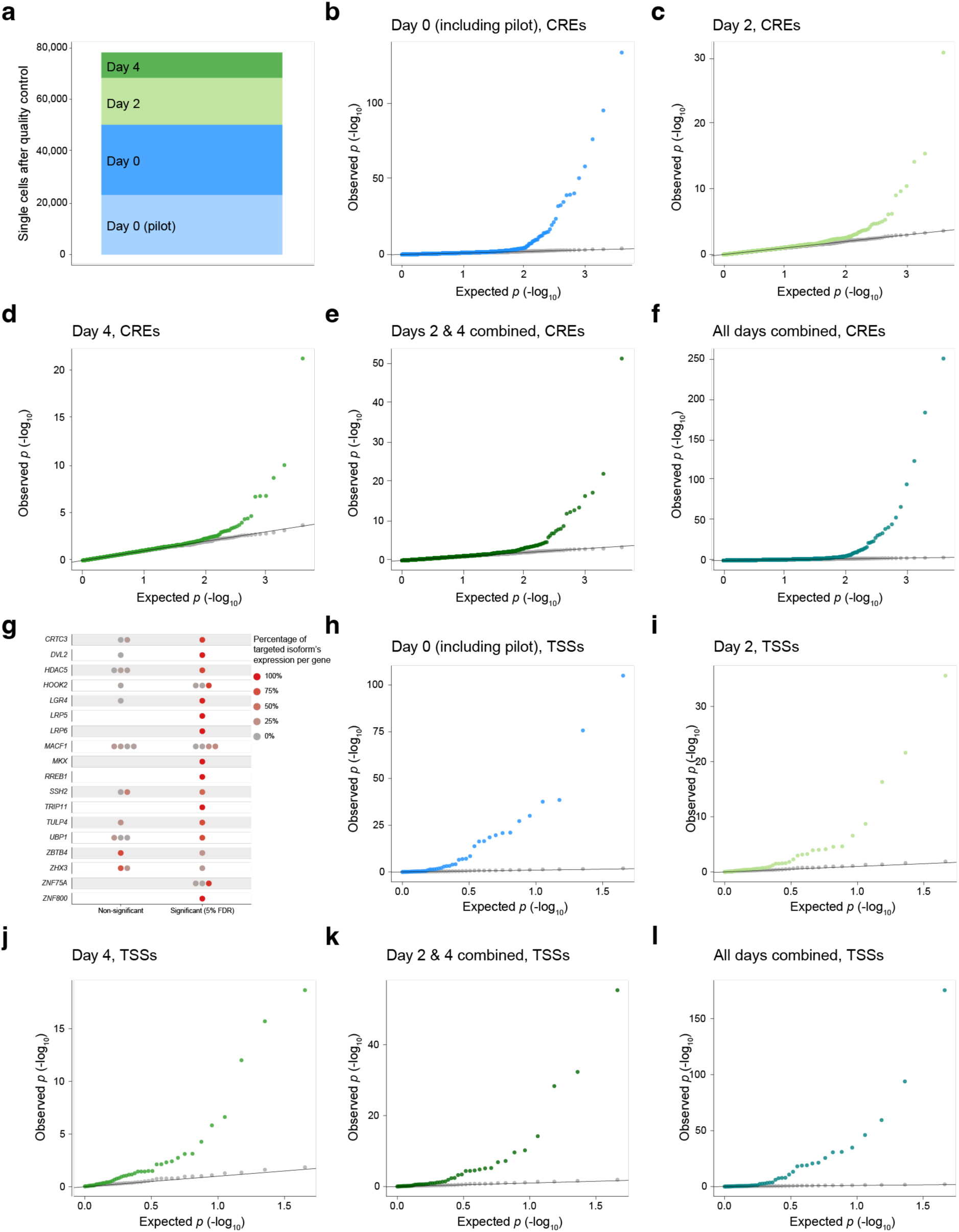
Summary of differential expression testing in cis for CRE- and TSS-targeting gRNAs. **a,** Total number of single cells available for differential expression testing after removing low quality cells and single cell multiplets. For both CRE- and TSS-targeting gRNAs, we performed differential expression testing for gRNAs targeting the same element against all genes in cis (within 1 Mb) and for non-targeting (NT) gRNAs and observed no NT–gene pairs significant at a 5% FDR across all conditions. **b–f,** Significant CRE–gene pairs (5% FDR) for gRNAs targeting the same cCRE, examining cells from Day 0 (**b**), Day 2 (**c**), Day 4 (**d**), Days 2 and 4 combined (**e**), and all time points combined (**f**). **g,** We assessed the effectiveness of gRNAs targeting gene TSSs and found that each targeted gene had at least one TSS whose inhibition produced a significant decrease in expression (5% FDR). **h–l**, Significant TSS–gene pairs (5% FDR) for gRNAs targeting the same TSS, examining cells from Day 0 (**h**), Day 2 (**i**), Day 4 (**j**), Days 2 and 4 combined (**k**), and all time points combined (**l**).

**Extended Data Fig. 5.**
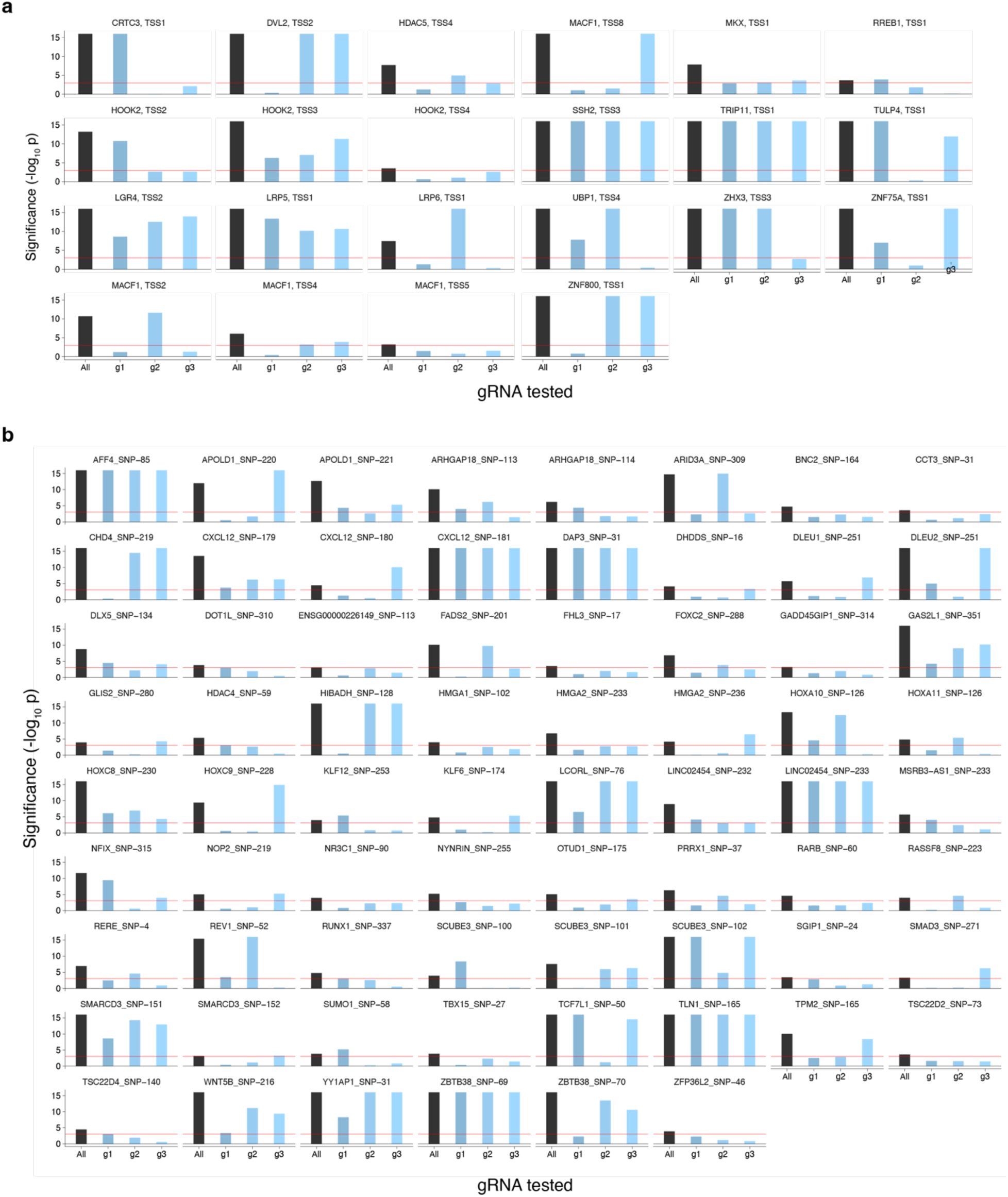
Individual gRNA effects for TSS- and CRE-targeting guides. Bars show the significance (-log_10_ *p*_SCEPTRE_) of the differential expression test using all three gRNAs combined ("All," navy blue bar) alongside each of the three individual gRNAs tested separately (g1, g2, g3; light blue bars). The red horizontal line marks the significance threshold corresponding to a 5% FDR cutoff. **a,** TSS-targeting gRNAs. Individual gRNAs largely reproduce the significance of the pooled test across target genes. **b,** CRE-targeting gRNAs. Most individual gRNAs independently reproduce the significance of the pooled test. For a subset of CRE–gene pairs, however, no single gRNA reaches significance on its own, and the association is detected only when combining all three gRNAs, reflecting the added statistical power of pooled testing to detect more modest regulatory effects.

**Extended Data Fig. 6.**
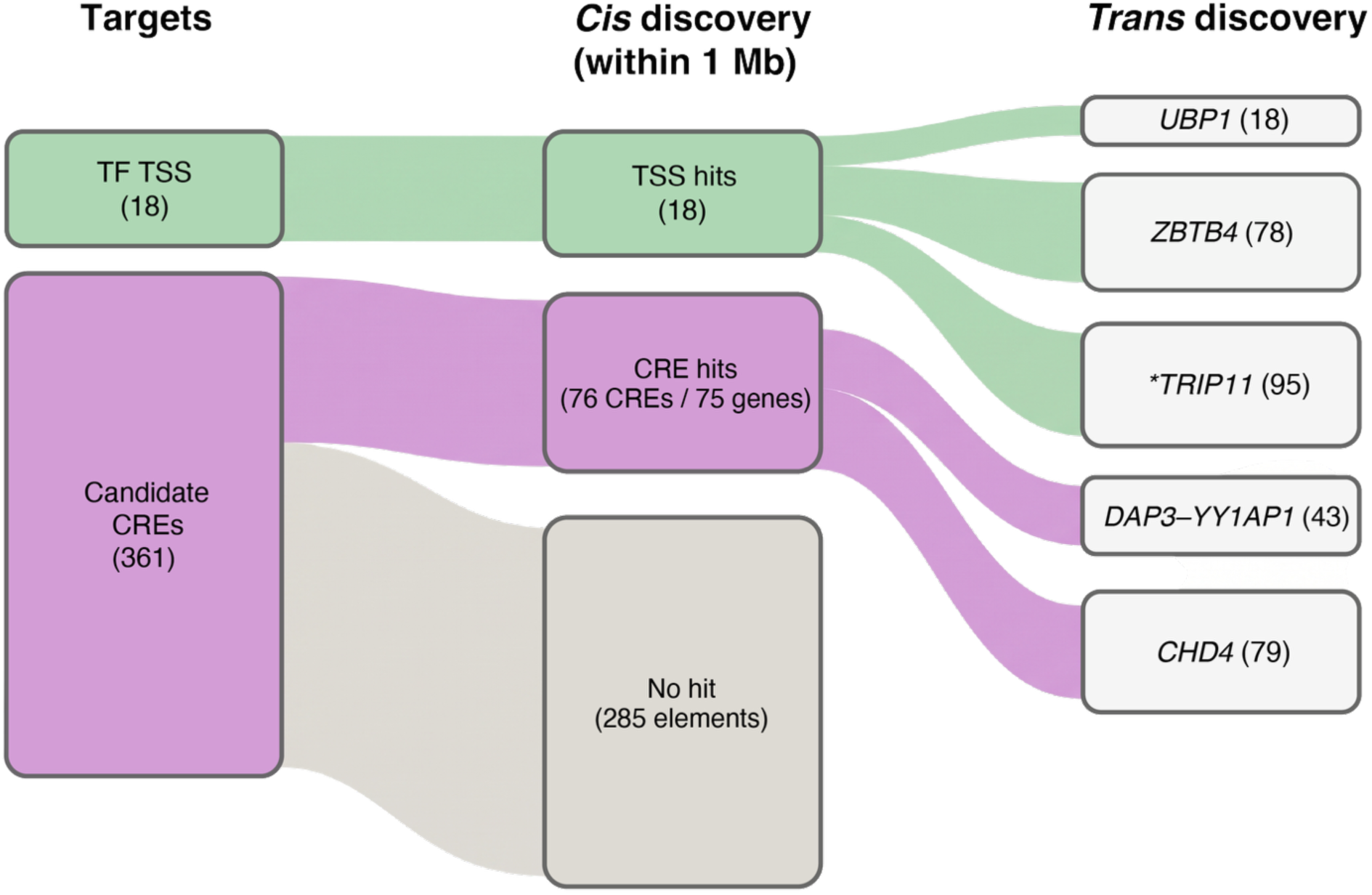
Summary of targeted elements and *cis* and *trans* discoveries in the STING-seq screen. Sankey diagram tracing the targeted elements (left) through *cis* discovery (middle) to *trans* discovery (right) in hFOB cells. * *TRIP11* was annotated as a potential TF but may not be a TF.

**Extended Data Fig. 7.**
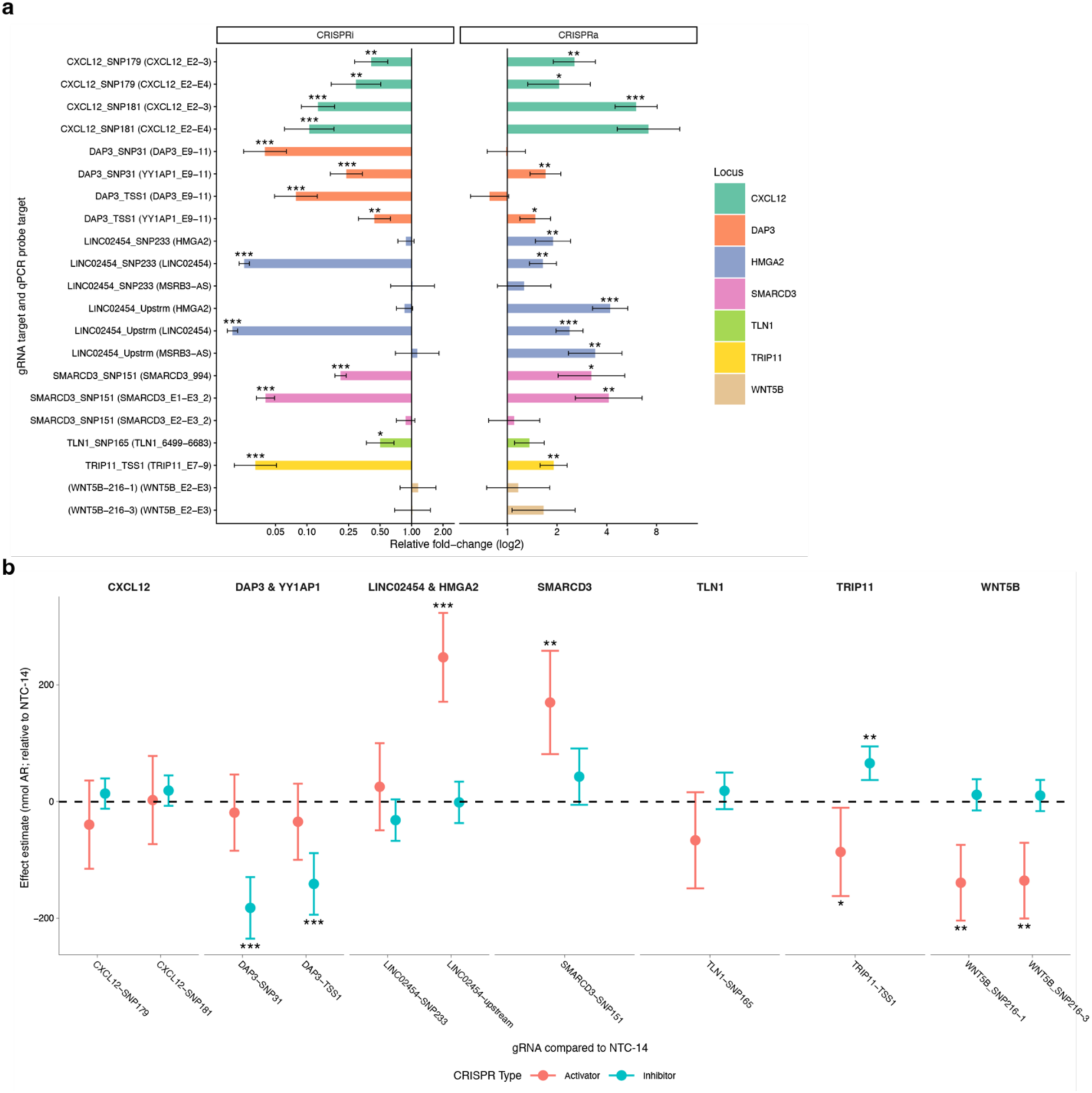
Arrayed CRISPRi and CRISPRa targeting of TSSs and CREs to measure changes in target gene expression and hFOB mineralization. We performed arrayed CRISPRi and CRISPRa lentiviral transductions in hFOB cells. **a,** Validation of the changes in gene expression observed with CRISPRi, and a test of whether the opposite effect is inducible with CRISPRa. **b,** Changes in mineralization measured by Alizarin red staining with CRISPRi and CRISPRa. Asterisks denote significant *p*-values from linear mixed models for arrayed experiments, comparing targets against non-targeting (NT) (* *p* < 0.05, ** *p* < 0.01, *** *p* < 0.001).

**Extended Data Fig. 8.**
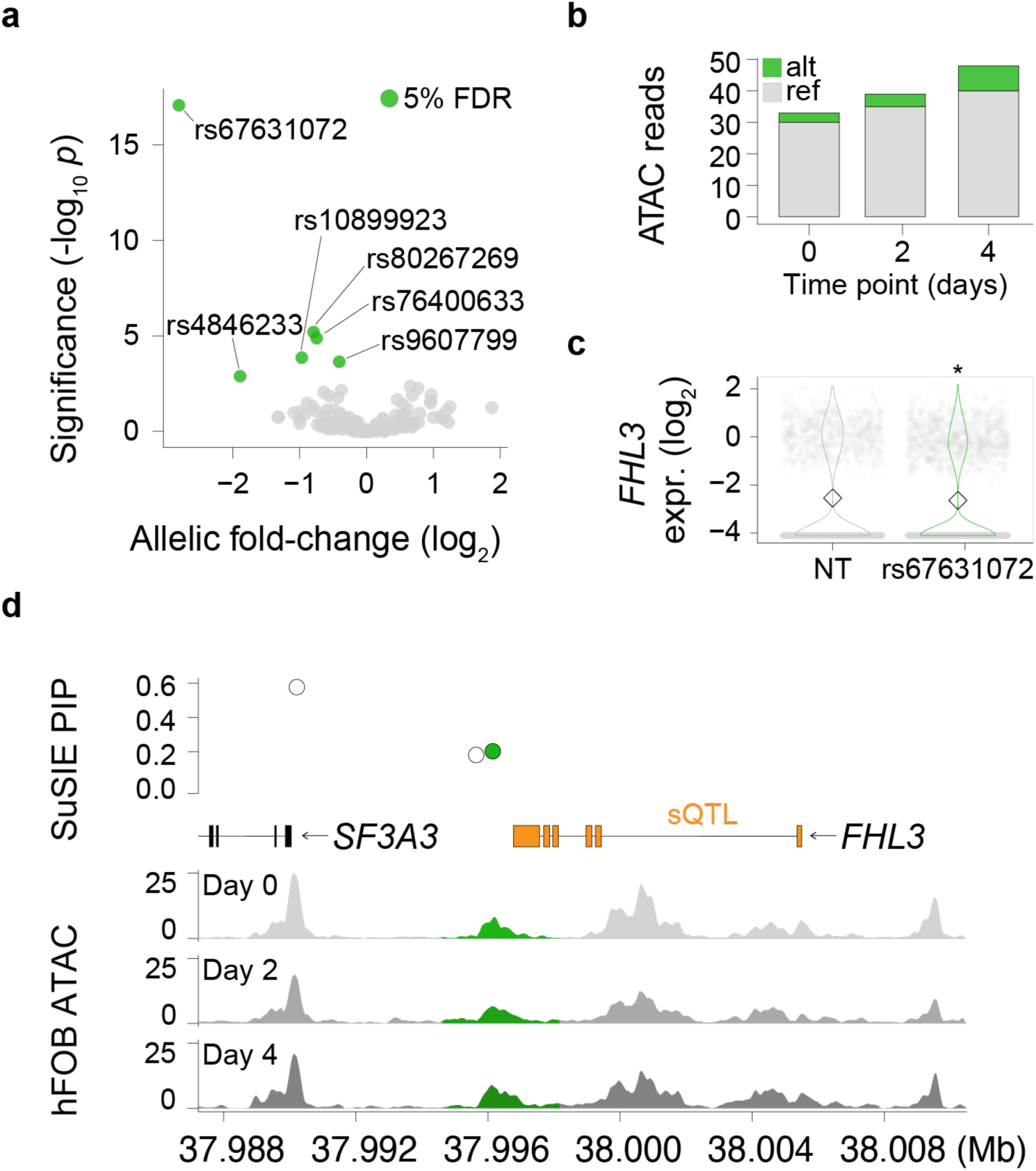
Allele-specific open chromatin in hFOB identifies a mechanism for the *FHL3* noncoding regulatory variant rs67631072. **a,** We performed allele-specific open chromatin analyses with hFOB ATAC-seq data and tested whether any targeted variants mapping to cCREs showed allele imbalance in ATAC-seq reads. **b,** We identified strong evidence of allele-specific open chromatin in hFOB cells for rs67631072. **c,** When we targeted the rs67631072-identified CRE in hFOB cells, we observed a significant decrease in *FHL3* expression (5% FDR). **d,** rs67631072 mapped downstream of the *FHL3* gene body and had previously been annotated as a splicing quantitative trait locus (sQTL) for *FHL3*. Asterisks denote *q*-values (Benjamini-Hochberg adjusted SCEPTRE *p*-values) significant at a 5% FDR.

**Extended Data Fig. 9.**
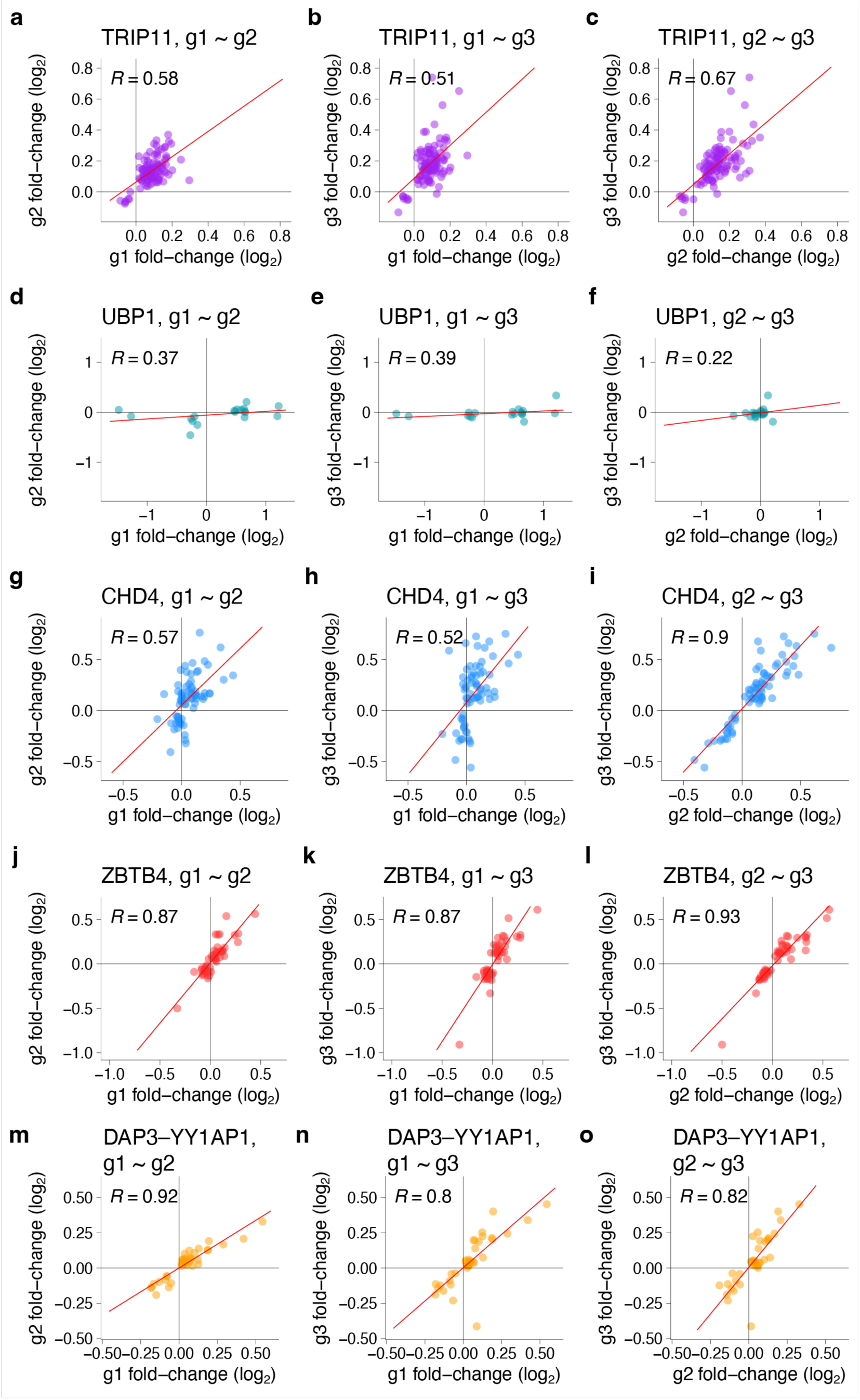
Guide-to-guide consistency of trans-regulatory effects identified by STING-seq. Pairwise comparisons of gene-level log_2_ fold-changes estimated from individual guide RNAs targeting *TRIP11* (**a–c**), *UBP1* (**d–f**), *CHD4* (**g–i**), *ZBTB4* (**j–l**), and *DAP3*–*YY1AP1* (**m–o**). Trans-regulated genes were defined using the pooled SCEPTRE analysis across all three guides. Each point represents a target gene, red lines indicate linear best fits, and Pearson correlation coefficients (*R*) are shown in each panel.

**Extended Data Fig. 10.**
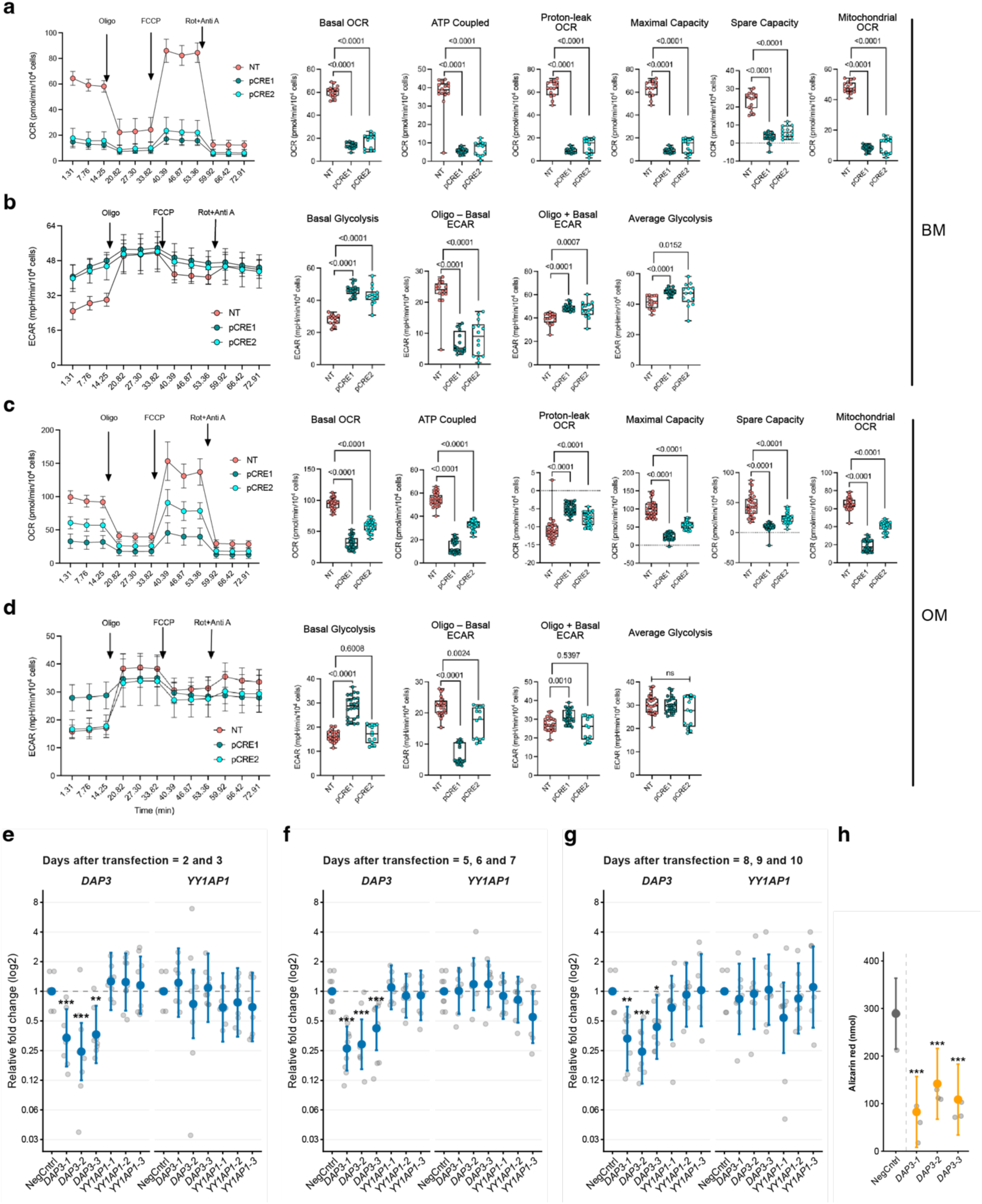
Mitochondrial function with *DAP3*–*YY1AP1* inhibition, and validation of siRNA knockdown and its effect on mineralization. **a–d,** Assays in basal and osteogenic media (BM and OM, respectively) to test mitochondrial function in hFOB cells with and without *DAP3*–*YY1AP1* inhibition. Under BM conditions we tested oxygen consumption rate (OCR) (**a**) and extracellular acidification rate (ECAR) (**b**) in hFOB cells transduced with *DAP3*– *YY1AP1* pCRE targeting and non-targeting (NT) gRNAs. We repeated these experiments under OM conditions, again testing OCR (**c**) and ECAR (**d**). **e–g,** Analyses of *DAP3* and *YY1AP1* gene expression by RT-qPCR, measuring transcript levels at days 2–3 (**e**), days 5–7 (**f**), and days 8– 10 (**g**) post-transfection. Displayed are the relative log_2_ fold-change of cells transfected with independent siRNAs targeting *DAP3* or *YY1AP1* transcripts relative to a non-targeting control (NegCntrl). **h,** Quantitative analysis of matrix mineralization via Alizarin red staining, transfecting cells with *DAP3* siRNAs and measuring calcium deposition relative to NegCntrl. Data in (**e–h**) are presented as model-estimated means with 95% confidence intervals (CI) derived from a linear mixed-effects model that accounts for inter-experiment variability. Individual data points represent normalized, independent biological replicates. Asterisks indicate statistical significance (* *p* < 0.05, ** *p* < 0.01, *** *p* < 0.001).

